# CYP3A5 inhibition causes G1/S blockade and synergizes with CDK4/6 inhibitor to suppress prostate cancer cell growth: Implications in reducing health disparity

**DOI:** 10.1101/2024.10.01.616176

**Authors:** Jeetesh Sharma, Imran K. Mohammed, Richard L. Tillett, Jake McLean, Shirley Shen, Ajay Singh, Oscar B. Goodman, Edwin C. Oh, Ranjana Mitra

## Abstract

Prostate cancer (PC) is a leading cause of death in men because of the high incidence and long-term inefficacy of the existing treatment options. Furthermore, it exhibits significant health disparities that affect African-American (AA) men more adversely than others do. Previously, we established CYP3A5, a highly expressed protein in AAs PC, as a positive regulator of androgen receptor (AR) signaling. We examined the impact of CYP3A5 depletion on genome-wide transcriptional output using RNA sequencing to gain deeper mechanistic insights. The data revealed that 561 genes were downregulated and 263 were upregulated upon silencing of *CYP3A5* in PC cells. Furthermore, *in silico* pathway analyses of differentially expressed genes suggested that the cell cycle regulation pathway was most significantly affected by *CYP3A5* inhibition. Cell cycle analysis of *CYP3A5*-silenced cells and those treated with clobetasol, a specific CYP3A5 pharmacological inhibitor, showed G1/S phase blockade. Both CYP3A5-depletion and pharmacological inhibition resulted in the downregulation of cyclin D, cyclin B, and CDK2, along with the upregulation of p27^kip1^ but had minimal effects on CDK4/6 levels. Combination treatment with clobetasol and the CDK4/6 inhibitor palbociclib exhibited synergy with combination index (CI) values ranging from 0.28-0.78. Our findings support the utility of CYP3A5 as a druggable therapeutic target that works more effectively in combination with CDK4/6 inhibition to limit the progression of PC, especially for AA patients with AA. This combination addresses CDK4/6 inhibitor resistance, which is often linked to CDK2 overexpression, and can potentially be useful in reducing disparities in the clinical outcomes of PC.

**Significance:** Our study highlights CYP3A5 as a key regulator of the cell cycle in prostate cancer (PC). Its overexpression in African American (AA) patients may be a key molecular driver of disparities in outcomes. The combination of CYP3A5 and CDK4/6 inhibitors shows a synergistic effect on therapeutic outcomes and addresses CDK2-mediated resistance. Thus, targeting both CYP3A5 and CDK4/6 could improve treatment outcomes, especially in AA PC patients.

## Introduction

Prostate cancer (PC) is the most prevalent malignancy and the second leading cause of cancer-specific mortality in American men [1]. African American (AA) men experience a disproportionate burden of PC with a 1.8 times higher incidence and more than twice the mortality rate than their non-Hispanic White American (NHWA) counterparts [2, 3]. Androgen signaling *via* the androgen receptor (AR) is a major regulator of PC growth, making androgen deprivation therapy (ADT) the primary treatment for locally advanced and metastatic PC [4]. However, despite the initial responses, ADT often fails, leading to the emergence of more aggressive castration-resistant PC (CRPC) [5]. Aberrant activation of AR signaling continues to remain significant for racial disparities in CRPC [6-8]. Understanding the underlying mechanisms driving these disparities, including the role of aberrant AR signaling, is crucial for developing more effective and equitable treatment strategies for African-American (AA) men with prostate cancer.

Several new systemic therapies have been deployed in clinics in recent years to treat advanced prostate cancer, such as darolutamide, apalutamide, and enzalutamide, which block AR transcriptional activity [9-11]; abiraterone, which blocks androgen biosynthesis [12]; and cabazitaxel, which is a tubulin-binding taxane [13]. However, patients often develop resistance to these modalities, limiting the survival of PC patients [14].

CYP3A5 plays a critical role in androgen receptor (AR) regulation by facilitating the nuclear translocation of AR, impacting the transcription of AR-dependent genes, and thus promoting PC growth [15-17]. CYP3A5, along with other members of the family, such as CYP3A4 and CYP3A43, is also involved in androgen metabolism by oxidizing testosterone to relatively less active hydroxy testosterone [18, 19]. However, unlike CYP3A4 and CYP3A43, CYP3A5 is expressed outside the liver, specifically in the prostate, and thus, can influence testosterone levels within the tumor microenvironment [20]. CYP3A5 is polymorphic, with 95% of non-Hispanic White American’s (NHWA) carrying the inactive *3 variant (A6986G, truncated protein) and 73% of AAs expressing the active *1 variant (wild-type), resulting heightened CYP3A5 expression in AAs [21, 22]. Additionally, CYP3A5 inducers or inhibitors can modulate intratumoral AR signaling, significantly affecting PC cell growth by either enhancing or suppressing AR-driven pathways [16]. However, its effects on other signaling pathways remain largely unexplored.

In this study, we sought to gain mechanistic insights into CYP3A5 functions by depleting its expression using siRNA in MDAPCa2b cells (AA-origin, expressing high levels of CYP3A5) followed by RNA sequencing. The data were later confirmed in a panel of other PC cell lines expressing different levels of the CYP3A5 protein. *In silico* analysis of differentially expressed genes identified the cell cycle regulatory pathway as the most affected, which was further validated by performing cell cycle analysis and immunoblotting of several cell cycle regulatory proteins. The treatment of PC cells with the highly selective CYP3A5 inhibitor clobetasol propionate [23], in combination with a CDK4/6 inhibitor (palbociclib), produced synergistic outcomes in all tested PC cell lines, including 22RV1, which expresses a constitutively active AR variant, ARV7. These data established CYP3A5 as a new druggable target in PC and demonstrated a mechanism that may be relevant for AA patients expressing elevated levels of CYP3A5 and exhibiting poor clinical outcomes.

## Material and methods

### Reagents

MTT (3-(4,5-Dimethylthiazol-2-yl)-2,5-Diphenyltetrazolium Bromide), DMSO (Dimethyl sulfoxide), trypsin (0.05%), antibiotic solution (100X) were purchased from Sigma-Aldrich Chemical Company, USA. Trypan blue 0.4 %, 1X DPBS (Dulbecco’s phosphate-buffered saline), and trypsin EDTA 0.5 % (1X) were procured from Gibco (USA). Protease (P8340) and phosphatase (524625) inhibitors were purchased from Millipore-Sigma (USA).

Primary antibodies against cyclin A2 (ab181591), Cyclin E2 (ab40890), Cyclin dependent kinase 2 (CDK2, ab32147), and CDK 6 (ab124821) were purchased from Abcam (USA). Cyclin D1, Cyclin B1, CDK4, p27 kip1, p-cdc2, and p-RB (ser807/811) were part of the sampler kit (9932T and 9917T), and along with p-Wee1 (4910T) were procured from Cell Signaling Technology (USA). GAPDH (10R-G109a) was used as the loading control (Fitzgerald, UK). Secondary antibodies (IR 680 and IR 800) were procured from LI-COR (Lincoln, NE).

### Cell lines and culture conditions

All the cell lines used in this study were purchased from ATCC. LNCaP, 22RV1, C4-2, and PC-3 cells were maintained in RPMI medium (Invitrogen, Carlsbad, CA, USA). MDAPCa2b and DU145 cells were maintained in F-12K medium (ATCC, USA) and IMEM (Invitrogen, Carlsbad, CA, USA), respectively. All the cell lines were supplemented according to ATCC guidelines and cultured under sterile conditions in a humidified 5% CO_2_ incubator at 37°C and were frequently evaluated for mycoplasma contamination using MycoAlert™ PLUS Mycoplasma Detection Kit (LT07-701) from Lonza, USA.

### Genotyping assay

CYP3A5 is located on chromosome 7q-21.1; the wild-type form is referred to as the *1 variant, whereas the most prevalent mutation, a single nucleotide polymorphism (SNP) at 6,986 A>G, is referred to as the *3 variant. A q-PCR-based genotyping method was used to detect allelic variants (*1 and *3) of CYP3A5. Genomic DNA was isolated from all cell lines using a QIAmp DNA mini kit (Cat No. 51304) from Qiagen according to the manufacturer’s protocol. A total of 10 ng genomic DNA per reaction was used for genotyping. To determine the allelic variants, we used the TaqMan Drug Metabolism Genotyping Assay (Cat no.4362691; CYP3A5*3 cat no. C_26201809_30), as previously described [16]. The genotyping assay was a probe-based predesigned assay (Applied Biosystems, USA). The VIC dye-labeled probe identified the wild-type (*1) allele, whereas the FAM dye-labeled probe identified the mutant (*3) allele. After the q-PCR run, allelic discrimination analysis was performed using 7500 Software v2.3 (Applied Biosystems) to detect the specific allele based on the amount of signal generated by reporter dyes (VIC and FAM).

### qRT-PCR

Total RNA was extracted using the RNeasy Mini Kit (Cat no.74104, Qiagen, Germantown, MD, USA), according to the manufacturer’s instructions. cDNAs were synthesized from 2.5 µg of total RNA using Qiagen’s RT2 first-strand kit (Cat No. 330404). The cDNA samples were diluted with 45 µl nuclease-free water and used as templates for qRT-PCR. The expression levels of CYP3A5 were quantified using an ABI 7500 Fast Real-Time PCR System with RT^2^ SYBR Green qPCR Master Mix (Cat No. 330502; Qiagen). Primers for CYP3A5 (ref position 73, NM_000777), spanning exon 1, were obtained from Qiagen (Cat. PPH01219F-200), and beta-2-microglobulin (B2M) were used as reference controls. Relative expression is presented as the fold-change of the target genes in treated cells compared to that in control cells.

### qRT-PCR based assay to determine CYP3A5 splice variants

We designed two sets of primers and probes (Table 1) to quantify the wild-type and mutant RNA-splice variants. The presence of the *3 SNP (mutant CYP3A5) results in a cryptic splice site, adding an extra 131bp (exon 3 B), which includes a stop codon. As a result, the CYP3A5*3 protein is truncated in homozygous (*3/*3) individuals with little active protein. We designed a FAM dye-containing probe at the junction of exon 3 and exon 4 to identify the wild-type (*1) variant (Table 1). The SUN dye-containing probe was designed against exon 3 B, which represents the mutant (*3) variant (Table 1). The probe and primer sets were custom synthesized from IDT and passed all quality checks using the probe primer analysis software. cDNA synthesis was performed using 2.3 µg of total RNA per 20 µl reaction. cDNA was further diluted with 45 µl nuclease-free water, and 3 µl of cDNA was used for qRT-PCR. Both forward and reverse primers were used at a concentration of 500nM each, whereas the probes (SUN*3/FAM*1) were used at a concentration of 200nM. The thermal cycler conditions used for PCR amplification included initial denaturation at 95°C for 5 min, followed by 40 cycles of denaturation (95°C,15 sec) and annealing/extension (60°C, 1 min).

**Table 1.**
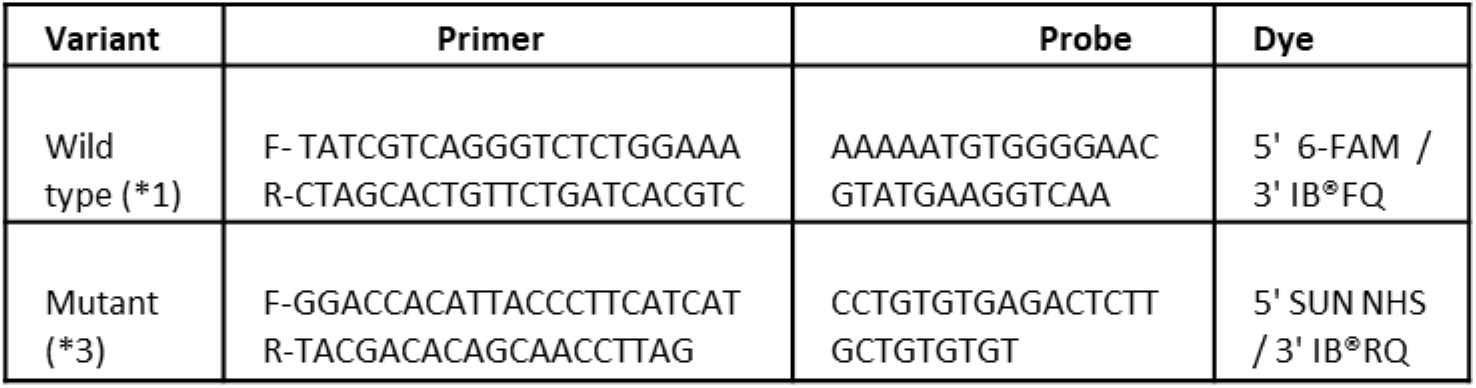
Primer and probe sequence used in splicing assay.

### siRNA transfection

Cells were seeded in complete medium on poly D-lysine-coated plates at a density of 80,000 cells per well for 6 wells and 2,000 cells per well in 96 well plates. Cells were transfected after 48 h using Lipofectamine RNAimax (Life Technologies, USA, Cat No. 13778-150) according to the manufacturer’s instructions. CYP3A5 expression was inhibited using a pool of four siRNAs (Dharmacon L-009684-01), and a smart pool non-target (NT) siRNA (Dharmacon D-001810-10) was used as a transfection control. The final concentration of the siRNA (NT and target) was 20 nM.

### Proliferation assessment by MTT assay

Cells were seeded in a poly D-lysine-coated 96-well plate (2,000 cells/well/100 µl) and transfected with siRNA (NT and target) 48 h after plating. Proliferation was assessed using the MTT assay (final conc. 400 µg/ml) 72 h post-transfection. MTT was added to each well and incubated for 2 h at 37°C. After incubation, the plates were centrifuged at 2,500 rpm for 10 min. The medium was discarded and 100 µl of DMSO was added per well. Absorbance was measured at 540 nm using a BioTek Synergy 2 microplate reader. The percentage cytotoxicity was calculated with respect to the non-target control cells. To evaluate the IC_50_ values GraphPad Prism software was used. Raw data were log-transformed and normalized using the software to generate cell growth figures.

### Methylene Blue assay

A methylene blue assay[24] was used to determine the effect of clobetasol and palbociclib on MDAPCa2b, LNCaP, and 22Rv1cells (IC_50_ and combination index). Methylene blue is a cationic stain that binds to DNA in the nucleus and RNA with low affinity in the cytoplasm. Cells were seeded (6,000-8,000 cells/well) in a poly D-lysine-coated 48-well plate to determine the CI index and 2,000-5,000 cells/well in a 96-well plate to determine IC_50_. After 48 h, the cells were treated with the drug alone or in combination at specified concentrations for 4 days (96 h). After the drug treatment cells were fixed with 10 % formalin in 1×PBS for 30 min. The fixative was removed, and cells were stained with filtered 1% methylene blue in 0.01 M borate buffer (pH 8.5). After 60 min of staining, the dye was removed, and the excess dye was washed (×3) with 0.01 M borate buffer. A mixture of ethanol and 0.1 M HCl (1:1, v/v) solution was added to dissolve the cell-bound methylene blue (100/200 µl per well for 96/48 well plates). The final absorbance was measured at 650 nm using a BioTek plate reader to determine the amount of methylene blue inside the cells. Methylene blue staining was used when the cells were treated with clobetasol, as it reacts with the MTT reagent to produce a color that could skew the growth assay results. The methylene blue assay was also used for the combinatorial treatment with palbociclib and clobetasol. A constant dose ratio of 20:1 (clobetasol: palbociclib) was used for combination studies and CalcuSyn software was used to calculate the combination index (CI) values.

### Clonogenic assay

Cells were seeded in a 6-well plate at a density of 2,000/4000 cells/well for LNCaP, 2,000 cells/well for 22RV1, and 1,000/2000 cells/well for MDAPCa2b. After 48 h, the cells were treated with clobetasol, palbociclib, or a combination of both, as previously described. Then, the cells were incubated at 37°C in a 5% CO_2_ incubator for two weeks. The culture medium was changed every 6 days using fresh drug. For staining, colonies were fixed (10% neutral buffer formalin) for 30 min, followed by staining with 0.01% crystal violet for 60 min, then washed, and detected using Odyssey CLx Imager. The assay was repeated three times.

### RNA sequencing and read analysis

For RNA sequencing, cells (40,000 cells/ml) were plated in 6 well plates and transfected as previously described. Total RNA was extracted 96 h post-transfection using a Qiagen RNeasy mini kit according to the manufacturer’s instructions. The RNA was quantified using a Nano Drop spectrophotometer. RNA integrity of each sample was assessed with an Agilent 2100 Bioanalyzer using RNA 6000 Nano chips (Agilent, Santa Clara, CA, USA) before library preparation and sequencing. RNA libraries were prepared from a total RNA input of 500 ng in 50 µl using an Illumina Stranded mRNA Prep Ligation kit according to the manufacturer’s instructions and sequenced using an Illumina NextSeq 500 instrument with a Mid Output kit (130M reads, 150 cycles). Sequencing was conducted at the GAA Core Lab of the Nevada Institute of Personalized Medicine at UNLV.

Illumina read pairs were trimmed to remove low-quality and synthetic adapter sequences using Trimmomatic version 0.39 [25]. Trimmed Illumina read pairs were aligned to the Ensembl reference human genome and transcriptome GRCh38 Release 104 [26] using the Hisat2 aligner, version 2.2.1 [27]. Transcripts observed per gene were summarized using Feature Counts software version 2.0.1 [28]. Differential expression between the treatments and controls was tested using the DESeq2 R package, version 1.30.1 [29], at a false discovery rate (FDR)-adjusted p-value of p_adj_ < 0.05. Sets of significant, differentially expressed genes were tested for shared membership in biochemical pathways, and Gene Ontology categories were inferred by over-representation analysis and gene set enrichment analysis using the GEne SeT AnaLysis Toolkit (WebGestalt) [30].

### Cell cycle assay

Cells (40,000 cells/well) were cultured in a 6-well plate followed by siRNA transfection or clobetasol treatment. After transfection/ treatment, cells were trypsinized and washed with 1× PBS. Cells were stained with PI in Nicoletti buffer (propidium iodide 50µg/ml, 0.1% sodium citrate, 0.1% Triton X-100, RNase 1 mg/ml, in 1×DPBS). Data were acquired using a C6 Accuri flow cytometer (Becton Dickinson, Mountain View, CA, USA) to study the effect of CYP3A5 inhibition on cell cycle arrest. Data were analyzed using the FlowJo software. The Dean-Jett-Fox model was used to calculate the percentage of the cell population at each cell cycle stage.

### Western blotting

Following transfection/ clobetasol treatment, cells were washed with 1×PBS (with 2 mM EGTA and 2 mM EDTA). RIPA buffer (50 mM TRIS, 150 mM NaCl, 1% NP40, 0.5% sodium deoxycholate, 0.1% SDS, 2 mM EDTA, 2 mM EGTA) supplemented with protease inhibitor and phosphatase inhibitor cocktail was added for lysis. The lysate was incubated for 20 min at 4 °C on an end-to-end rotor followed by centrifugation at 16,000 ×g for 20 min at 4 °C. The supernatant was collected in a separate tube, and the protein concentration was estimated using a Micro BCA™ Protein Assay Kit (cat no. 23235) from ThermoScientific, USA. An equal amount of protein was loaded for SDS-PAGE in 4-12 % BoltTM Bis-Tris premade gel from Invitrogen for 80 min at 120 V and blotted on to a nitrocellulose membrane. The membrane was blocked with 5% BSA in 1×TBST, followed by overnight incubation with primary antibodies. A fluorescence-labeled secondary antibody was used for detection using an Odyssey CLx Imager. GAPDH was used as a loading control.

### Data Availability

RNA sequencing data were deposited using GSE xxxxx. All other data are available as results, supplementary data, or figures.

## Results

### Prostate cancer cells of AA origin express high levels of functional CYP3A5

We performed genotyping and expression analysis of CYP3A5 in a panel of six prostate cancer cell lines, MDAPCa2b, LNCaP, 22RV1, DU145, C4-2, PC-3, and MDAPCa2b. Genotyping data revealed that MDAPCa2b (AA origin) carried one wild-type*1 allele (*1/*3), whereas all other NHWA origin lines carried both mutant alleles (*3/*3) of CYP3A5 (Figure 1 A, Figure S1) [16]. Similarly, MDAPCa2b cells also showed higher *CYP3A5* expression than the other NHWA cell lines (Figure 1 B).

**Figure 1.**
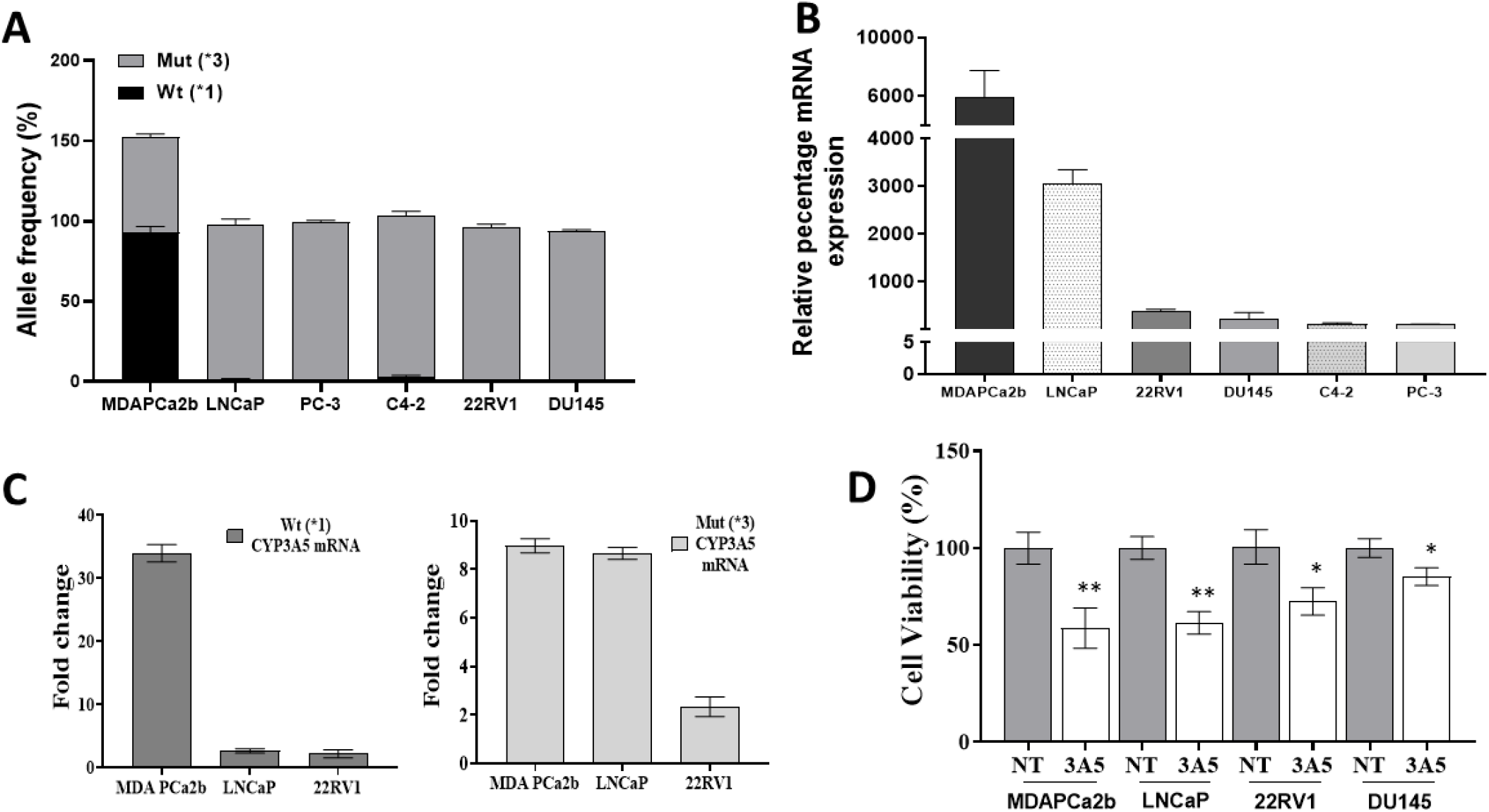
CYP3A5 mRNA expression in prostate cancer (PC) cell lines and effect of CYP3A5 siRNA on PC cell growth. **(A)** qPCR-based genotype assay was used to determine the presence of wild-type (*1) or mutant (*3) CYP3A5 polymorphism in six PC cell lines as described in the methods. **(B)** Quantification of total CYP3A5 mRNA expression in the PC cell lines was done using SYBR green-based qRT-PCR assay. Total RNA (2.3 µg) in a reaction mixes of 20 µl was used for cDNA synthesis. Beta2-microglobulin was used as an internal reference control. The graph represents the mean ± SD of three independent experiments. **(C)** CYP3A5 splice variants (*1-wild type, generating full-length protein and *3-mutant, carrying extra exon 3B containing stop signal resulting in truncated protein) were quantified in mentioned PC cell lines using specifically designed primer-probe based qRT-PCR assay. The separate primer and probe set were used to detect the variants as described in Table (1). **(D)** CYP3A5 inhibition blocks the growth of PC cells. Cells were plated in a 96-well plate followed by transfection with CYP3A5 siRNA and NT siRNA pool (non-target control). The graph represents the mean ± SD of three independent experiments MTT assay was performed, and absorbance was taken at 540 nm after 96 hours to assess the cell growth. Data represents the mean ± SD of three independent experiments performed in quadruplets. *P*-value was calculated with respect to NT control. *P-*value * ≤ 0.05, ** ≤ 0.005

Although LNCaP carries both mutant variants (*3/*3), our results showed a higher expression of *CYP3A5* mRNA in the LNCaP line (Figure 1B). This result could be due to the detection of both mutant and wild-type mRNA transcripts using our primer set (located in exon one). To test whether the higher *CYP3A5* mRNA expression observed in LNCaP was due to the location of the primer set, we designed another probe/primer set that could differentiate between wild-type and mutant transcripts (containing the exon 3 B region with a stop codon), as described in the Methods section (Table 1). MDAPCa2b expressed both a wild-type and mutant transcript in line with its genetic makeup, and the other two lines, LNCaP and 22RV1, expressed the mutant mRNA transcript (Figure 1C). LNCaP and 22RV1 NHWA cell lines carrying (*3/*3) mutation also expressed 5-10% wild-type transcript, possibly because they can bypass the splicing events to some extent, as described previously [22, 31]. Although LNCaP cells express high total *CYP3A5* mRNA (Figure 1B) compared to other NHWA PC lines, only a fraction of transcripts are correctly spliced (Figure 1C) and can produce full-length active CYP3A5 protein [17]. Furthermore, depletion of CYP3A5 with a pool of four siRNAs led to reduced growth in all four tested PCa cell lines (MDAPCa2b, LNCaP, 22RV1, and DU145). The pool of CYP3A5 siRNAs blocked 70-80% of the *CYP3A5* mRNA expression and downregulated CYP3A5 protein levels by 4-5-fold, as shown previously [15, 16]. highest growth inhibition (~40%) was observed in MDAPCa2b and LNCaP cells, followed by 22Rv1 (~30%), and DU145 (~10%) (Figure 1D).

### CYP3A5 depletion affects gene expression associated with important biological pathways

To identify the signaling downstream of CYP3A5, we performed RNA-seq on MDAPCa2b cells (AA origin, CYP3A5 overexpressing) that had been subjected to CYP3A5 knockdown using a pool of siRNAs, along with their corresponding NT (non-target) siRNA control. Differential expression analysis using the DSeq2 R package revealed 1814 differentially expressed genes (DEGs) (Table S1) with an adjusted p-value of ≤ 0.05 and LFC (limit fold change) shrinkage applied to log2 fold changes (Figure 2A). A heat map generated after applying further stringent conditions (genes with log2 fold change of two or more, and p-value of ≤ 0.01) showed 649 DEGs (Figure 2B), of which 187 were upregulated and 462 were downregulated in the CYP3A5 knockdown cells. Kyoto Encyclopedia of Genes and Genomes (KEGG) pathway analysis with DEGs showed several putative molecular pathways either upregulated (2) or downregulated (11) in *CYP3A5* silenced cells with a false discovery rate (FDR) of ≤ 0.05 (Figure 2C). Several of these downregulated genes were involved in growth regulatory pathways, including cell cycle progression (39 genes) (Table 2), cellular senescence (16 genes), DNA replication (18 genes), and base excision repair (9 genes) (Figure S2, Table S2), which supported our previous observation of growth reduction with *CYP3A5* siRNA treatment.

**Table 2.** Effect of CYP3A5 siRNA treatment on cell cycle regulatory genes in MDAPCa2b cell line.

| Genes | Fold Change | Adj *p*-value | Genes | Fold Change | Adj *p*-value |
| --- | --- | --- | --- | --- | --- |
| CDK2 | -4.30918 | 1.43E-21 | MAD2L2 | -2.34728 | 0.004015 |
| CCNE1 | -2.6974 | 3.22E-10 | BUB1B | -2.84183 | 5.03E-05 |
| CCNA2 | -3.3294 | 2.17E-09 | BUB1 | -2.99189 | 8.88E-08 |
| CDK1 | -3.23554 | 3.55E-12 | CDC20 | -3.27981 | 6.05E-14 |
| CCNB2 | -3.46486 | 3.96E-22 | CDC25B | -2.07827 | 4.01E-11 |
| CCNB1 | -3.44683 | 4.94E-30 | YWHAH | -2.59293 | 1.32E-22 |
| PLK1 | -3.42183 | 2.28E-36 | CDC25A | -2.5158 | 9.04E-13 |
| CDC25C | -2.9214 | 2.07E-10 | PCNA | -2.21665 | 1.93E-10 |
| E2F1 | -2.62421 | 2.31E-09 | PKMYT1 | -3.21612 | 1.01E-12 |
| CDC6 | -3.29623 | 9.13E-24 | ORC1 | -2.50341 | 7.03E-12 |
| CDC45 | -2.95471 | 5.04E-16 | ORC6 | -2.20295 | 2.89E-11 |
| MCM2 | -2.64697 | 2.20E-15 | E2F2 | -2.49838 | 9.14E-12 |
| RBL1 | -2.73966 | 7.12E-06 | CDC7 | -1.92477 | 2.12E-04 |
| SKP2 | -2.59813 | 1.94E-07 | DBF4 | -1.87302 | 0.045855 |
| CHEK1 | -3.53091 | 3.27E-11 | MCM7 | -1.84184 | 1.00E-05 |
| CHEK2 | -2.03068 | 2.65E-08 | MCM3 | -1.97495 | 6.38E-14 |
| TTK | -2.73307 | 0.001764 | E2F6 | -1.58026 | 0.003628 |
| ESPL1 | -2.32616 | 1.26E-04 | E2F5 | 1.567193 | 0.019534 |
| PTTG1 | -3.64238 | 8.90E-08 |  |  |  |

**Figure 2.**
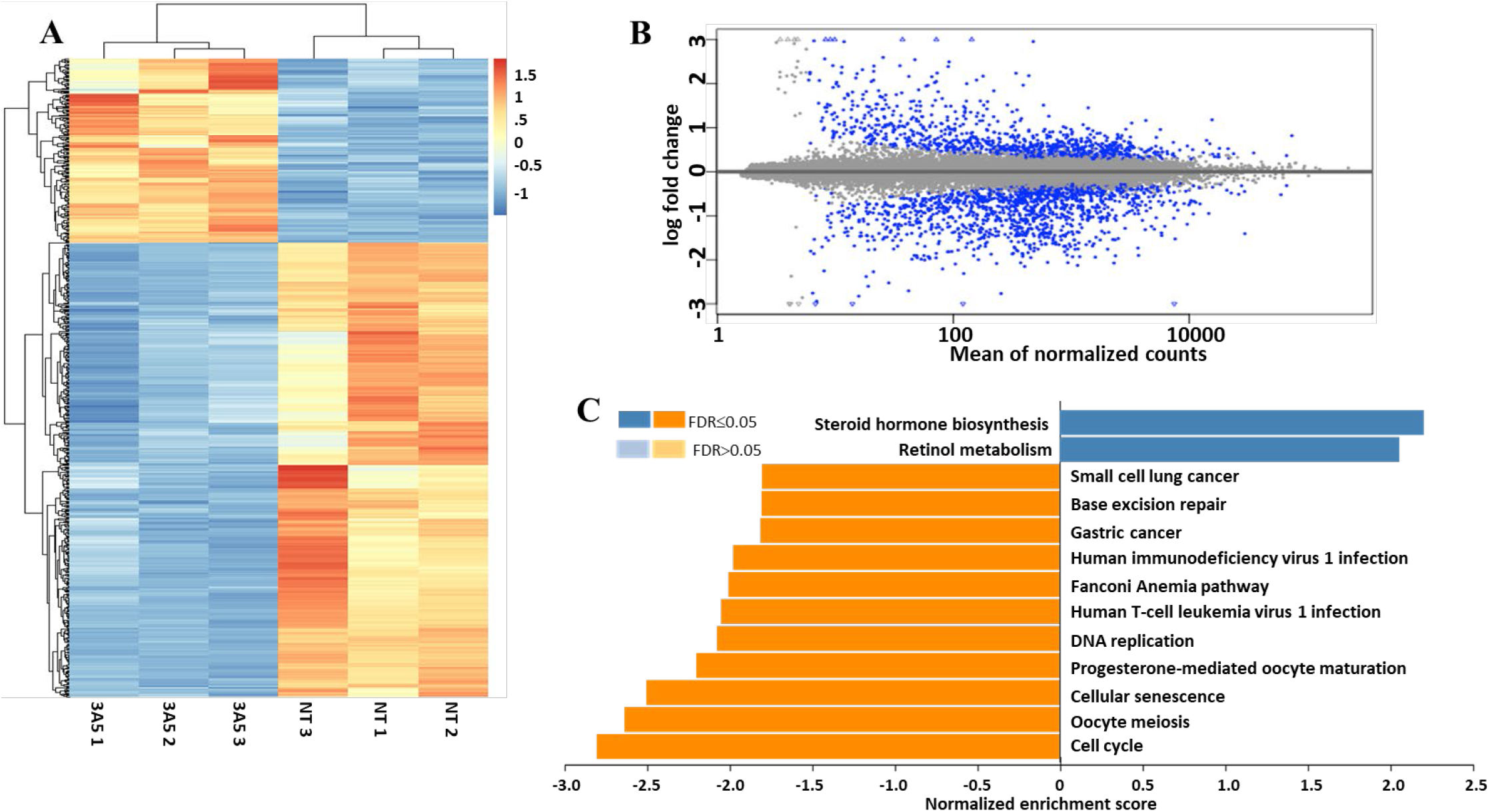
RNA sequencing studies show that CYP3A5 siRNA treatment downregulates the expression of several growth regulatory genes. RNA sequencing was performed with RNA isolated from MDAPCa2b cells, 96 hours after transfection with NT (non-target) and CYP3A5 siRNA as described in the methods. (A) Hierarchical cluster heat map showing the differential expression of genes between the NT and CYP3A5 siRNA treated MDAPCa2b cells. Genes that displayed ≥log2-fold difference in expression were identified and plotted for the heat map. A total of 649 significantly (p-adj <0.01) differentially expressed genes relative expression across the triplicate of two samples (CYP3A5 target vs non-target siRNA) is shown. Orange shows higher expression and blue shows lower expression. (B) Distribution of expression level (x-axis) and fold-change magnitude (y-axis) in RNA-seq of CYP3A5-treated *vs*. non-target treated cells. Genes with significant changes in expression are indicated in blue, those with no significant changes in expression are in grey. (C) Normalized gene set enrichment analysis (GSEA) of differentially expressed genes identified several pathways either up or downregulated with a false discovery rate (FDR) of ≤0.05 with the Kyoto Encyclopedia of Genes and Genomes (KEGG) pathway database. Cell cycle pathway was the most significantly downregulated (37 genes) pathway detected by the pathway analysis.

The cell cycle pathway was the most significantly affected pathway, with 39 genes involved in this pathway being downregulated (Table 2). Most downregulated genes were specific to the cell cycle phase, whereas some had overlapping roles in two or more phases of the cell cycle (Figure S2A). We observed transcriptional downregulation of several cyclins and their specific kinase partners, cyclinE/CDK2 (G1 phase), cyclinA/CDK2 (S phase), cyclinA/CDK1, and cyclinB/CDK1 (G2 phase) with CYP3A5 siRNA treatment. Of the other downregulated mRNAs, p107 (RBL1-Rb like protein), E2F 1,2 (cyclin E transcriptionskp2 (regulating p27 function), ckh1,2 (cdc25A and G2/M phase regulator), PLK1 (cdc25C and G2/M phase regulator), and Myt1 (G2/M transition) are all known cell cycle regulators. The expression of the cdc25 family, cdc25A/B/C phosphatase, was also downregulated by CYP3A5 siRNA knockdown. In addition, we observed downregulation of several genes, including p107 and p130, Rb-like proteins that are part of the dream complex (inhibit the G1/S transition), cell division cyclin genes (cdc-6,7,20,45), origin complex recognition genes (orc-1,6), and mini-chromosome maintenance (mcm-2,3,7), which are known to regulate cell cycle progression.

Among the downregulated genes in the cellular senescence pathway, some targets overlapped with the cell cycle regulatory pathway, such as cyclin A2, cyclin B1/B2, cyclin E1, CDK1/2, E2F1/2, CHEK1, CDC25A, and RBL1, whereas others were specific to senescence pathways, such as mitogen-activated protein kinase 6 (MAP2K6), forkhead box M1 (FOXM1), calmodulin-like 5(CALML5), and lin-9 DREAM MuvB core complex component (LIN9) (Figure S2B, Table S2). Most of the genes downregulated in the DNA replication pathway are part of the replication complex: DNA polymerase α-primase complex, DNA polymerase d complex, DNA polymerase e complex, mini chromosome maintenance complex (mcm 2-7), replication protein A3, clamp (PCNA), clamp loader (RFC2/4 and RFC3/5), Fenl, and DNA ligase (Lig1) (Figure S2D, Table S2). The base excision repair pathway shared some downregulated genes similar to those in the DNA replication pathway (POLD1/3, POLE2, LIG1, PCNA, and FEN1), whereas NEIL3, PARP2, and HMGB1 were specific to the repair pathway (Figure S2C, Table S2). In addition, several other pathways were affected, as identified by KEGG analysis (oocyte meiosis, oocyte maturation, T-cell leukemia virus infection, Fanconi anemia, human immunodeficiency virus infection, gastric cancer, small cell lung cancer, retinol metabolism, and steroid hormone biosynthesis). Although some of these are not directly related to cell growth, the same common genes as mentioned above play a significant role in regulating these pathways (Figure S2E-F, Table S2). In small cell lung cancer and gastric cancer pathways, Myc downregulation can reduce the expression of CKS1 and Skp2, which participate in the degradation of p27, an inhibitor of CDK2/cyclin E complex formation, thereby causing G1/S phase cell cycle arrest. In addition, downregulation of the Wnt-frizzled pathway, Ras, and c-Met, which play a vital role in cancer growth and metastasis, with CYP3A5 siRNA treatment.

### CYP3A5 inhibition by RNA interference causes cell cycle arrest at G1/S check point

Prostate cancer cell lines (LNCaP, MDAPCa2b, 22RV1, and DU145) were treated with an siRNA pool targeting CYP3A5, along with a non-target (NT) control pool, for 96 h and evaluated for their effect on each cell cycle phase.

The cell cycle assay was performed as described in the Methods section and analyzed using the Dean-Jett-Fox (DJF) univariate method (Figure 3). All cell lines showed a significant accumulation of cells in the G0/G1 phase. LNCaP, MDaPCa2b, and 22RV1 showed maximum accumulation of cells in the G0/G1 phase, followed by DU145, compared to NT control siRNA-treated cells (Table 3). We also observed a significant reduction in the S-phase cell population in all the cell lines.

**Table 3.** CYP3A5 siRNA causes G1/S block in PC cells.

|  |  | G0/G1 | S | G2/M |
| --- | --- | --- | --- | --- |
| MDAPCa2b | NT siRNA | 70.9±2.1 | 20.0±0.7 | 7.7±3.5 |
|  | 3A5 siRNA | 79.1±2.2 | 5.7±4.1 | 15.3±1.8 |
| LNCaP | NT siRNA | 60.8±1.8 | 19.5±5.6 | 19.8±3.8 |
|  | 3A5 siRNA | 77.0±0.7 | 7.6±2.9 | 15.6±3.0 |
| 22RV1 | NT siRNA | 53.6±2.5 | 31.5±1.1 | 14.2±2.5 |
|  | 3A5 siRNA | 71.0±1.1 | 23.9±2.5 | 4.2±4.2 |
| DU145 | NT siRNA | 34.3±0.4 | 34.9±0.5 | 27.7±0.2 |
|  | 3A5 siRNA | 39.7±2.8 | 31.3±0.9 | 25.0±2.2 |

**Figure 3.**
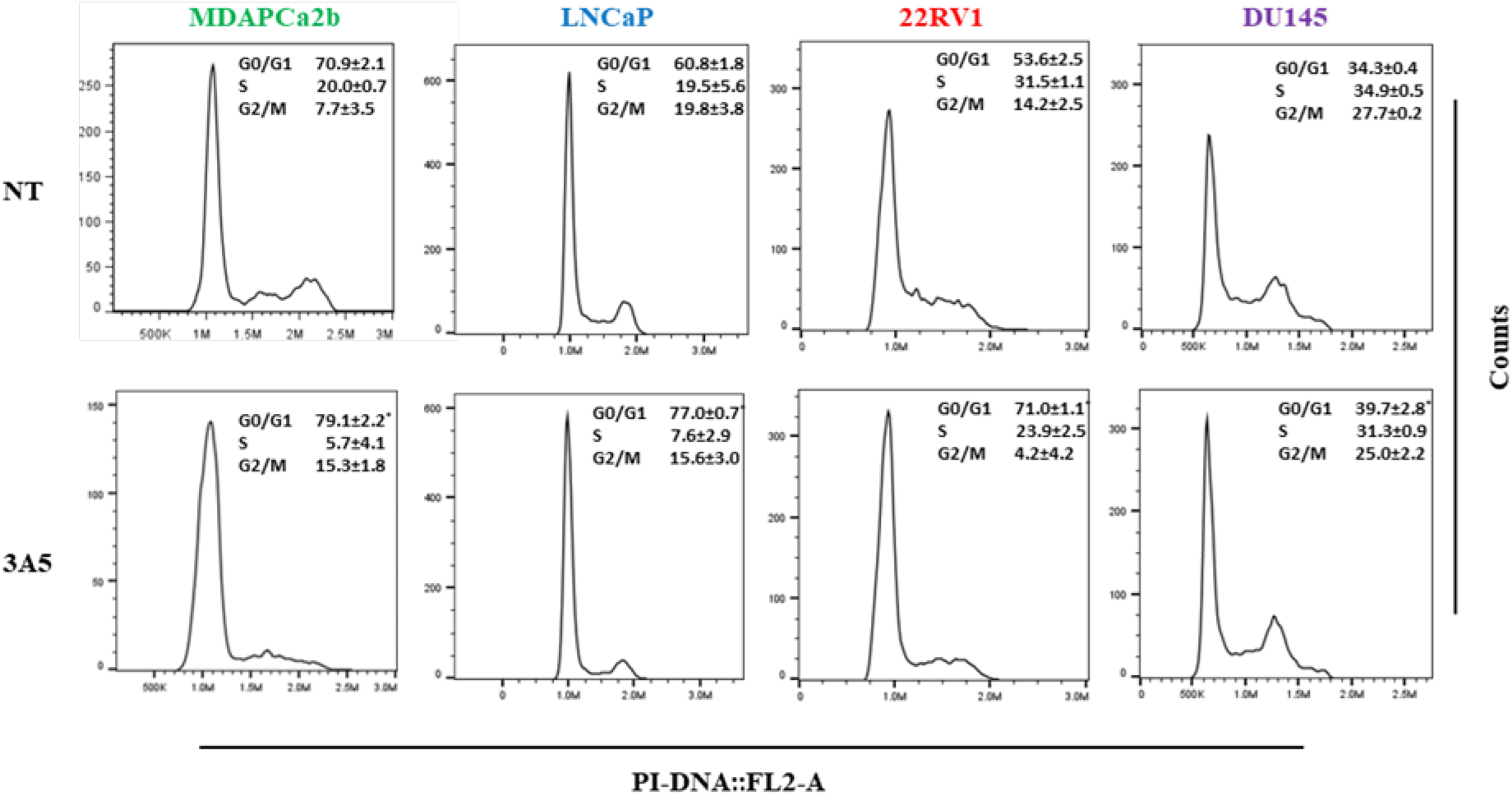
Inhibition of CYP3A5 using siRNA causes accumulation of G0\G1 cells. Dean Jett Fox cell cycle analysis showing the frequency of G0/G1, S, and G2/M population in non-target control (NT), and CYP3A5 siRNA treated cells. Cell cycle was performed 96 hours after treatment with NT and CYP3A5 siRNA with propidium iodide staining in Nicolette buffer. The experiments were repeated three times, and the mean value for all the experiments are shown for each population. Two-way ANOVA was employed to determine the significant change in cell cycle phases in NT and CYP3A5 siRNA treated sets \**p*≤0.05.

Western blot analysis was performed to further evaluate the differential levels of cell cycle regulatory proteins in MDAPCa2b, LNCaP, and 22RV1 lines (Figure 4). As we observed an increase in cells in the G0/G1 phase, we examined the levels of G0/G1 cyclins and their corresponding kinases (cyclinD/CDK4/6 and cyclinE/CDK2).

**Figure 4.**
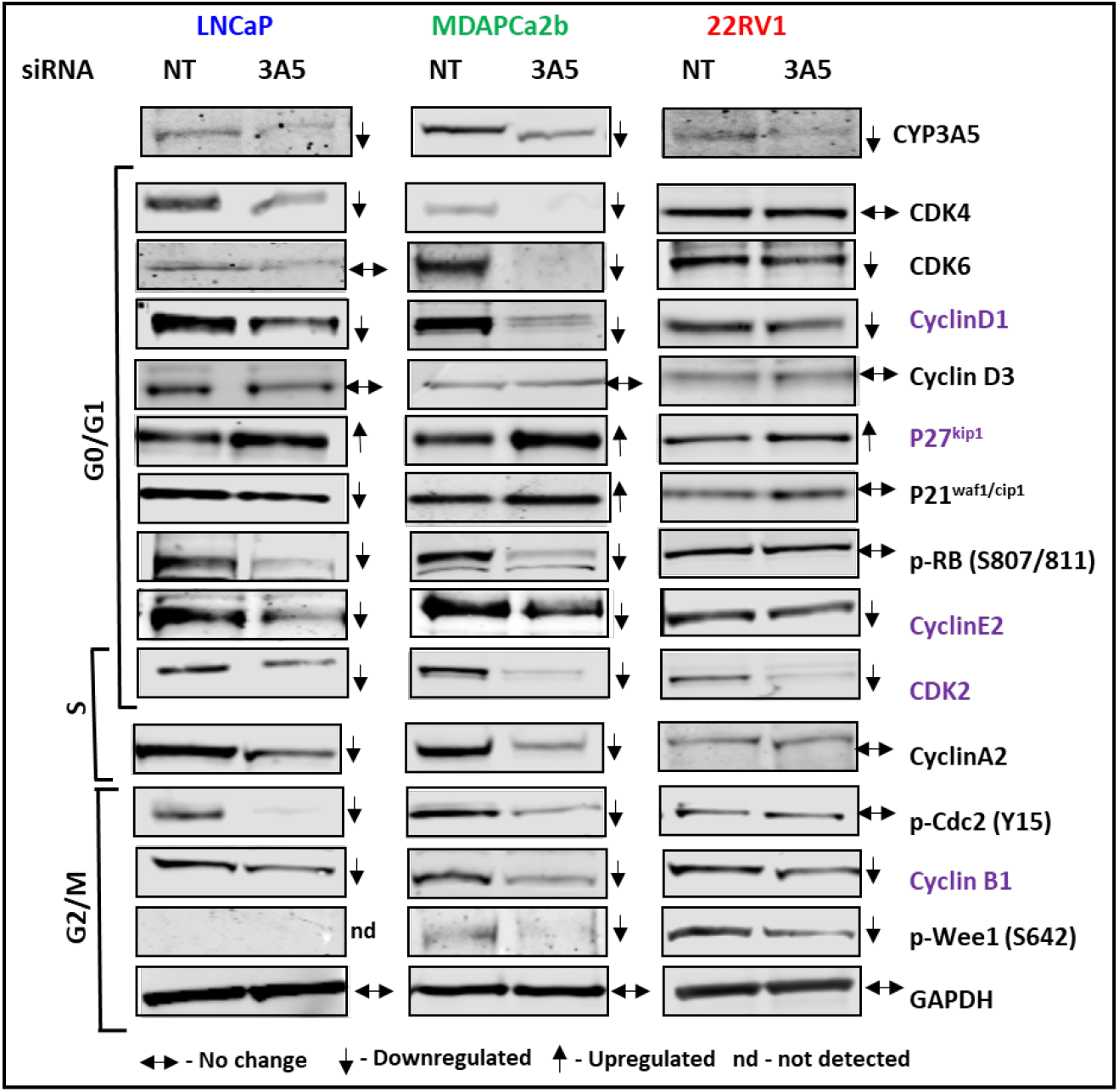
Differential regulation of cell cycle regulating protein after CYP3A5 inhibition. Western blot analysis exhibited the involvement of cell cycle regulatory protein in siRNA-treated PC cell lines (MDAPCa2b, LNCaP, and 22RV1) compared to the non-target (NT) control siRNA pool (96 h). GAPDH was taken as a loading control. The experiment was repeated at least twice; a representative replicate is shown.

Given the potential block in the G1/S transition, we observed a reduction in cyclin D1 in all three cell lines evaluated (LNCaP, MDAPCa2b, and 22RV1) and downregulation of CDK4 in LNCaP and MDAPCa2b, whereas CDK6 downregulated MDAPCa2b and 22RV1 after CYP3A5 siRNA treatment. Cylin D3 levels were not affected in any of the evaluated lines. The other two negative regulators in the G1/S phase were p21^cip1^ and p27^kip1^; p21^cip1^ levels increased only in MDAPCa2b cells, and p27^kip1^ levels increased in all three cell lines, suggesting that CYP3A5 modulates its expression to regulate the G1/S transition. The levels of INK p16/p18 regulators either did not change or were undetectable (data not shown) in any of the lines tested. The late G1 phase cyclin, cyclin E, was downregulated in LNCaP and MDAPCa2b cells, but not in 22RV1 cells, although the corresponding kinase, CDK2, was downregulated in all three lines. Increased Rb phosphorylation is known to directly upregulate cyclin E transcription, which explains the downregulation of Rb phosphorylation observed only in LNCaP and MDAPCa2b cells, and not in 22RV1 cells.

The S phase cyclin A levels were downregulated in LNCaP and MDAPCa2b cells, but did not change in 22RV1 cells. CDK2, which pairs with Cyclin A in S phase, was downregulated in all three lines. G2/M cyclin B expression was downregulated in all three cell lines after CYP3A5 siRNA treatment. G2/M entry is regulated by five nodes: PLK1, Chk1, Cdc25c, Wee1, and CDK1. Depending on cellular stress levels, these nodes determine the phosphorylation state of CDK1 (Cdc-2) at Y15. Our results indicate that Y15 phosphorylation is downregulated upon CYP3A5 inhibition, which renders the cyclin D/CDK1 complex inactive (LNCaP and MDAPCa2b cells). Y15 phosphorylation of CDK1 is often dependent on the phosphorylation of Wee-1 at S642, which is downregulated in MDAPCa2b and 22RV1 cells. In the absence of cellular stress, CDK1 can transfer phosphate (Y15) to Cdc25c, activating the CyclinB/CDK1 complex and proceeding to mitosis [32]. All fold changes shown in Table 4 were examined after normalization to the corresponding GAPDH levels (Figure 4).

**Table 4.** Fold change in cell cycle regulatory protein after CYP3A5 inhibition.

| Proteins | LNCaP |  | MDAPCa2b |  | 22RV1 |  |
| --- | --- | --- | --- | --- | --- | --- |
|  | NT | 3A5 | NT | 3A5 | NT | 3A5 |
| CDK4 | 100 | 56 | 100 | 66 | 100 | 101<sup>NC</sup> |
| CDK6 | 100 | 88<sup>NC</sup> | 100 | 10 | 100 | 65 |
| CyclinD1 | 100 | 58 | 100 | 15 | 100 | 76 |
| CyclinD3 | 100 | 93<sup>NC</sup> | 100 | 92<sup>NC</sup> | 100 | 103<sup>NC</sup> |
| CDK2 | 100 | 69 | 100 | 12 | 100 | 32 |
| Cyclin E1 | 100 | 99<sup>NC</sup> | 100 | 85 | 100 | 94<sup>NC</sup> |
| Cyclin E2 | 100 | 32 | 100 | 32 | 100 | 83 |
| pRB ser 807/811 | 100 | 25 | 100 | 15 | 100 | 99<sup>NC</sup> |
| p27 | 100 | 150 | 100 | 167 | 100 | 136 |
| p21 | 100 | 63 | 100 | 131 | 100 | 103<sup>NC</sup> |
| Cyclin A2 | 100 | 30 | 100 | 73 | 100 | 99<sup>NC</sup> |
| Cyclin B1 | 100 | 56 | 100 | 53 | 100 | 74 |
| p-cdc2 | 100 | 17 | 100 | 28 | 100 | 85<sup>NC</sup> |
| p-Wee1 | 100 | ND | 100 | 58 | 100 | 62 |

The table represents the normalized fold changes after GAPDH calibration and is representative of Figure 4.
ND- Not detected, NC- (No change, P>0.05 mean all replicate experiments)

### Clobetasol, a selective CYP3A5 inhibitor, suppresses prostate cancer cell growth and induces G1/S arrest, mirroring the effects of CYP3A5 siRNA

Our study established that the inhibition of CYP3A5 negatively regulates prostate cancer growth by affecting the levels of cell cycle regulatory proteins. To corroborate these findings, we evaluated clobetasol, the only commercially available CYP3A5 inhibitor [23]. We used MDAPCa2b, LNCaP, and 22RV1 cells as model cell lines and determined their IC_50_ values using a methylene blue-based growth assay. The IC_50_ doses for the three lines were: MDAPCa2b 10.46 µM, LNCaP 19.89 µM and 22RV1 12.88 µM based on results from three replicates (Figures 5A, 5D). Interestingly, we observed lower IC_50_ values for clobetasol using the clonogenic assay in all three cell lines (LNCaP-1.16µM, MDAPCa2b-0.83µM, and 22RV1 1.74µM) (Figure 5B, 5D), suggesting its use in translational studies at lower dosages with fewer side effects. To confirm that the inhibition of PC growth by clobetasol was not due to its steroid activity, we treated PC cells with prednisone (another steroid). Prednisone-treated PC cells showed growth inhibition only at high concentrations (Figure 4E), and the predicted IC_50_ (~76-100µM) were all above the highest dose of prednisone (67.5µM) used in the assay.

**Figure 5.**
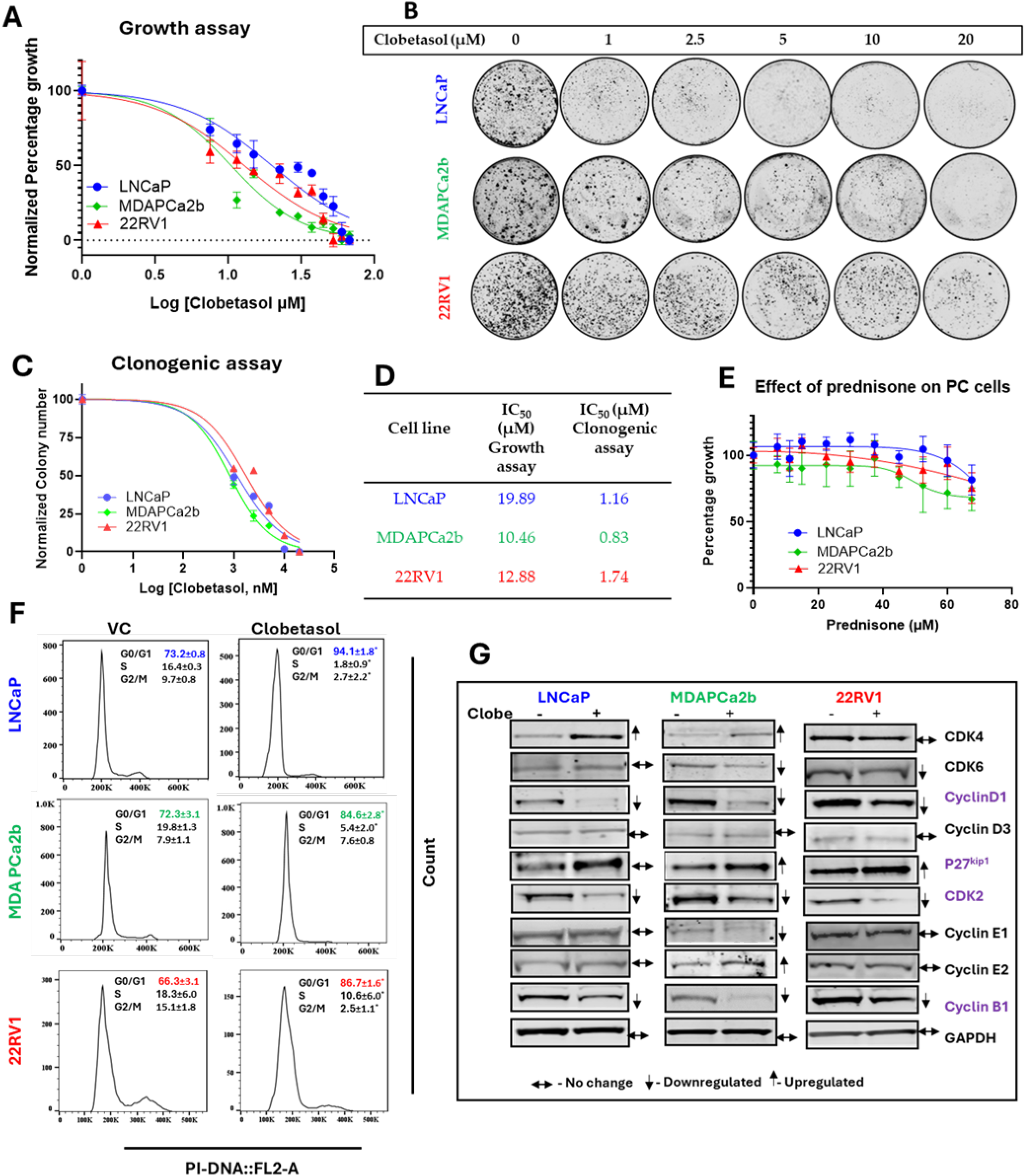
Clobetasol a specific CYP3A5 inhibitor blocks the growth of PC cells and causes G1/S block. **(A)** Cells were plated in a 96-well plate and were treated with an increasing dose of clobetasol (7.5 µM to 67.5 µM) after 48 hours of plating. Cell growth was determined using methylene blue staining at 96 hrs. after treatment. The experiment was repeated three times, each experiment was done in quadruplicate. The plot represents one of the replicates and was plotted after log transformation and normalization using GraphPad Prism software. **(B)** Clonogenic assay was performed by plating either 2000 (22RV1) or 4000 (MDAPCa2b and LNCaP) cells per well in a 6 well plate and treated with clobetasol (1-20µM) for two weeks. Colonies were then fixed and stained with crystal violet as described in methods. **(C)** The plot represents the percentage number of colonies after treatment with different dosage of clobetasol as shown in 5B. The graph was plotted using GraphPad Prism software after counting the actual number of colonies in each set (triplicate), normalization and log transformation. **(D)** The IC_50_ values were calculated using GraphPad Prism for the methylene blue (growth assay) and the clonogenic assays. **(E)** Cells were plated in a 96-well plate and were treated with an increasing dose of prednisone (7.5 µM to 67.5 µM) after 48 hours of plating. Cell growth was determined using MTT assay at 96 hrs. after treatment. The experiment was repeated three times, and the graph plotted using GraphPad Prism is a representative experiment. **(F)** As mentioned, PC cells were treated with clobetasol (20 µM), and cell cycle analysis was performed at 48 hrs. Dean-Jett-Fox cell cycle analysis was conducted in FloJo and a representative histogram plot is shown. The experiment was conducted in replicates (minimum of three) and ordinary Two-way ANOVA was employed to determine the significant change in cell cycle phases in treated versus vehicle control (VC-DMSO). \**p*≤0.05. **(G)** Western blot analysis exhibiting the effect of clobetasol (20uM) treatment on PC cell lines (96 h). GAPDH was taken as a loading control. The experiment was repeated a minimum of two times, a representative replicate is shown.

Cell cycle data analysis after post-clobetasol treatment using the DJF method revealed accumulation of cells in the G0/G1 phase in all the three lines tested: MDAPCa2b (84.6±2.8), LNCaP (94.1±1.8) and 22RV1 (86.7±1.6) compared to vehicle control treated MDAPCa2b (72.3±3.1), LNCaP (73.2±0.8) and 22RV1(66.3±3.1) (Figure 5F). We also found a significant reduction in the number of cells in S-phase in all three lines. In addition, we observed a reduction in the G2/M cell population in LNCaP and 22RV1 cell lines but not in the MDAPCa2b cell line. Taken together, our cell cycle data for clobetasol-treated cells were in line with the cell cycle data for CYP3A5-siRNA-treated cells, and both treatments led to cell cycle arrest at the G1/S checkpoint.

Western blotting to determine the effect of clobetasol on cell cycle regulatory proteins revealed similar effects to CYP3A5 siRNA treatment. Cyclin D1, cyclin B1 and CDK 2 were downregulated, and p27 was upregulated in all three PC cell lines evaluated following clobetasol treatment (Figure 5G, Table 5). Interestingly, CDK4 was upregulated in LNCaP and MDAPCa2b cells and downregulated in 22RV1 cells treated with clobetasol. In contrast, CDK6 levels remained unchanged in LNCaP cells but were downregulated in MDAPCa2b and 22RV1 cells.

**Table 5.** Fold change in cell cycle regulatory protein after Clobetasol treatment.

| Proteins | LNCaP |  | MDAPCa2b |  | 22RV1 |  |
| --- | --- | --- | --- | --- | --- | --- |
|  | NT | 3A5 | NT | 3A5 | NT | 3A5 |
| CDK4 | 100 | 195 | 100 | 186 | 100 | 85 |
| CDK6 | 100 | 98<sup>NC</sup> | 100 | 20 | 100 | 64 |
| CyclinD1 | 100 | 34 | 100 | 20 | 100 | 42 |
| CyclinD3 | 100 | 110 | 100 | 93<sup>NC</sup> | 100 | 98<sup>NC</sup> |
| CDK2 | 100 | 32 | 100 | 20 | 100 | 53 |
| CyclinE1 | 100 | 102<sup>NC</sup> | 100 | 41 | 100 | 108 |
| CyclinE2 | 100 | 106 | 100 | 240 | 100 | 96<sup>NC</sup> |
| p27 | 100 | 158 | 100 | 176 | 100 | 198 |
| CyclinB1 | 100 | 76 | 100 | 56 | 100 | 74 |

### Combination treatment with clobetasol and palbociclib (CDK4/6 inhibitor) synergistically inhibited the growth of PC cells

Cell cycle is often dysregulated in cancer cells and 20-40% of PCs show an upregulation of CDK4/6 and an abrogation of Rb-1(retinoblastoma) negative regulation. The CDK4/6 inhibitors palbociclib, ribociclib, and abemaciclib are currently in clinical trials for PC. Since these CDK4/6 inhibitors require positive expression of Rb, their use is restricted to only Rb1-expressing tumors [33-35]. Our experiments showed that the primary target of CYP3A5 inhibition in all tested cell lines was CDK2 (both with siRNA and clobetasol). Therefore, we wanted to test the chemical inhibition of CYP3A5 (using clobetasol) in combination with the CDK4/6 inhibitor (palbociclib) to increase its applicability to a broader population. Additionally, clobetasol treatment upregulated CDK4 in MDAPCa2b and LNCaP cells (Figure 5G), making them a good target for CDK4 inhibitor combination studies. To evaluate the possible synergy between these two drugs, we determined the IC_50_ value of palbociclib (CDK4/6 inhibitor) in the three lines. Cells were treated with palbociclib (0.0125 µM to 15.6 µM) for 96 h, and the methylene blue assay was used to determine the growth and IC_50_, which ranged between (0.3-0.6 µM). To further investigate the synergy between clobetasol and palbociclib in MDAPCa2b and LNCaP cells, the cells were treated with clobetasol and palbociclib, alone or in combination, at a constant dose ratio of 20:1 (clobetasol: palbociclib) for 96 h (Figure 6B, C). The Combination index (CI) was calculated after measuring the growth using methylene blue staining. CalcuSyn software was used to calculate the CI index at four different dosages, and a simulation was used to calculate the ED_50_ and ED_75_ (effective dose) (Table 6). We found that the combined treatment of clobetasol and palbociclib was more effective in MDAPCa2b (ED_50_, CI=0.025) than in LNCaP (ED_50_, CI=0.253), as was also relevant by the CI at different dose combinations (lower the CI – higher synergy). As we observed that clobetasol and palbociclib synergistically reduced prostate cancer cell growth, we further confirmed the long-term growth inhibitory effect of both drugs, alone and in combination, using a clonogenic survival assay. In all three cell lines, MDAPCa2b, LNCaP, and 22RV1 colonies were significantly reduced when treated with a combination of clobetasol and palbociclib compared to when treated with either drug alone (Figure 6D).

**Table 6.**
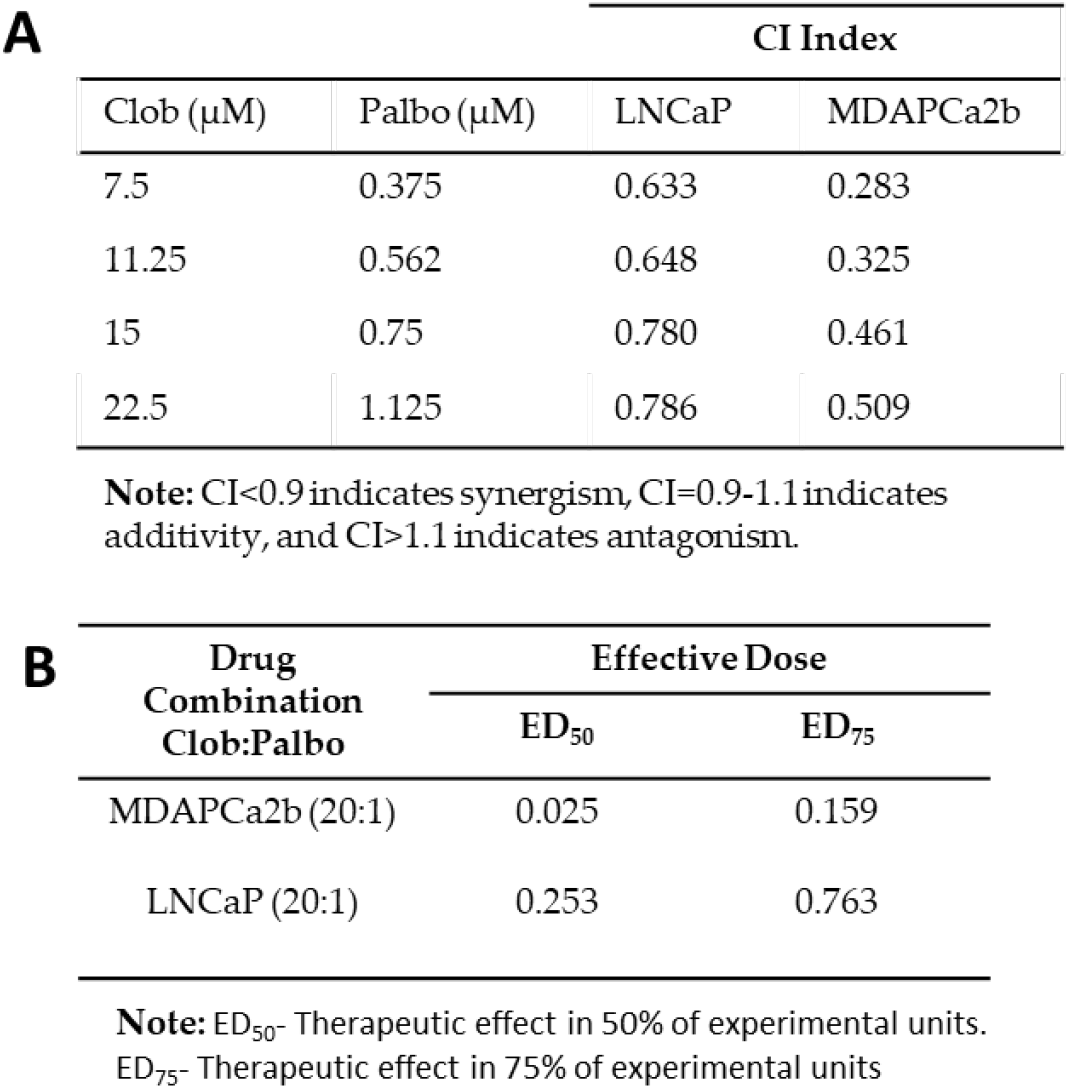
Calculated CI value at an effective dose (ED_50_ and ED_75_) after treating MDAPCa2b and LNCaP in a constant ratio of clobetasol (Clob) and palbociclib (Palbo).

**Figure 6.**
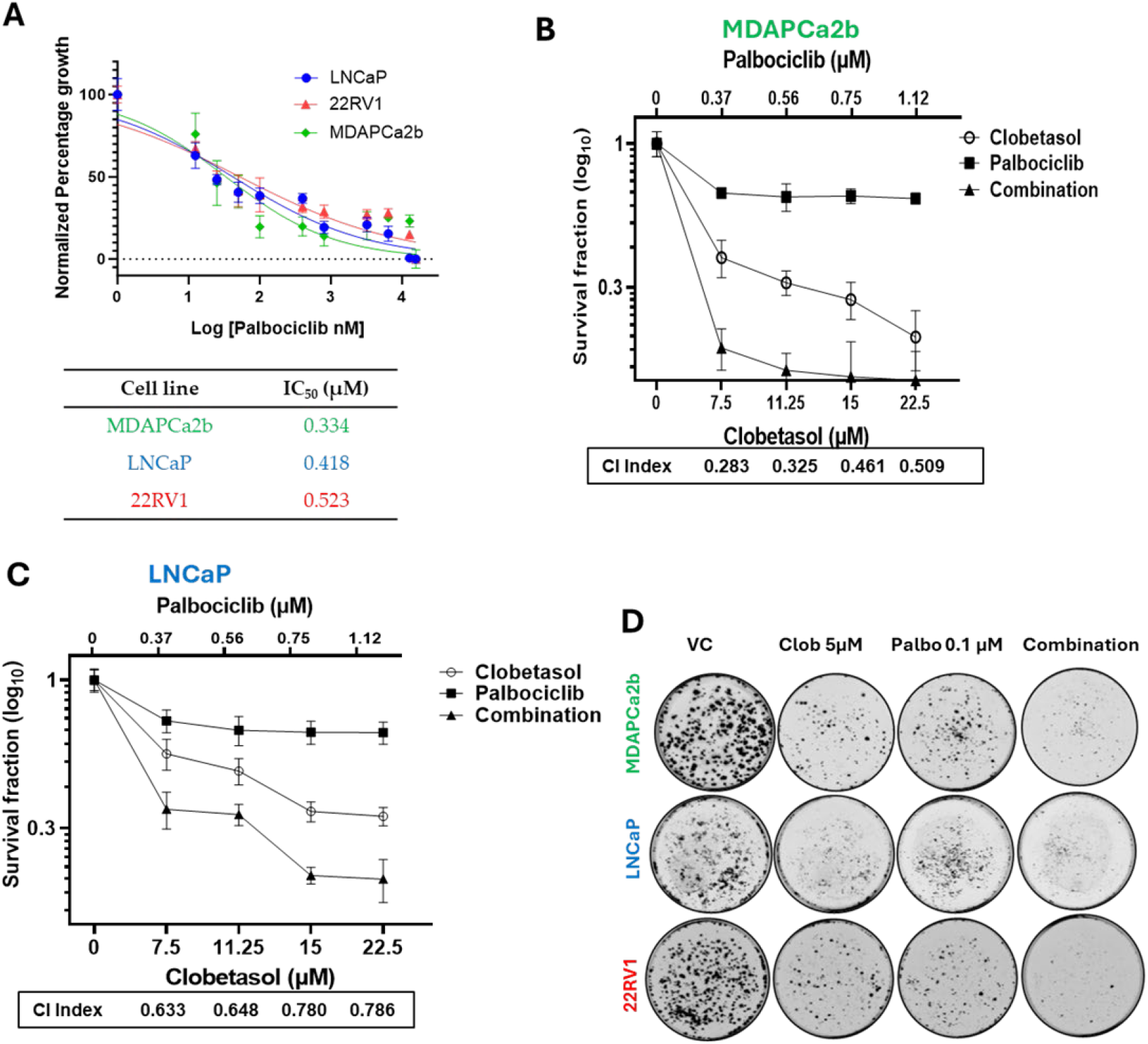
Cotreatment with clobetasol and Palbociclib synergistically decreases cell growth of prostate cancer cells. **(A)** Prostate cancer cells were treated with increasing doses of palbociclib (0.0125 µM to 15.6 µM) and cell growth was determined using methylene blue at 96 hrs. The experiment was repeated thrice in quadruplets; the graph shown is representative. The IC_50_ values were calculated using GraphPad Prism (interpolation method). **(B&C)** Cells were treated at indicated doses with clobetasol and Palbociclib with a constant ratio of (20:1), individually or in combination, and cytotoxicity was measured by methylene blue assay after 96 hours. The combination index (CI) shown in the graph was calculated using CalcuSyn software for each drug concentration. Experiments were repeated three times, and a representative plot is shown. **(D)** Clonogenic assay showing the number of colonies formed after 2 weeks of treatment with clobetasol and palbociclib individually or in combination at the indicated dose. Colonies were fixed and stained with crystal violet dye for visualization. The experiment was performed three times, and a representative experiment is shown.

## Discussion

In this study, we have uncovered significant insights into the role of CYP3A5 in regulating prostate cancer cell growth. Previous research from our laboratory established that inhibiting CYP3A5 leads to a reduction in prostate cancer growth [15-17]. Moreover, we demonstrated that CYP3A5 plays a pivotal role in promoting nuclear translocation and activation of AR, which is crucial for prostate cancer growth.

CYP3A5 is a polymorphic gene with 12 known loss-of-function SNPs, most of which result in a nonfunctional protein [21]. Notably, these SNPs exhibit a high degree of ethnic-specific linkage disequilibrium. The most common SNP, referred to as *3, results from a point mutation at 6986 (A>G), leading to a nonfunctional truncated protein. This mutation is present in 95% of non-Hispanic white Americans (NHWA), and 75% of African Americans (AAs) carry the wild-type gene, referred to as *1. Our genotyping of several PC cell lines confirmed that the *3 mutation is prevalent in NHWA; however, because of the limited availability of AA-origin cell lines, we could only test MDAPCa2b, which confirmed the presence of one pair of wild-type and one pair of mutant (*1/*3) CYP3A5 variant. Our investigation revealed that correctly spliced CYP3A5 mRNA, resulting in an active protein, is overexpressed in MDAPCa2b cells. In LNCaP and 22RV1, some correctly spliced CYP3A5 mRNA was also present, possibly because the SNP was bypassed during mRNA processing. Typically, 5-10% of correctly processed RNA are observed in cryptic splice variants [22].

Our RNA sequencing data underscores the importance of CYP3A5 in regulating cell growth by revealing a decrease in the mRNA expression of several genes involved in cell cycle regulation, cellular senescence, and DNA damage repair pathways that are often dysregulated in cancer cells. In particular, the cell cycle regulatory pathway stood out, with the mRNA expression of 37 genes being downregulated upon CYP3A5 siRNA treatment. Cell cycle progression relies on the coordinated activation of cyclin-dependent kinases (CDKs) paired with their respective cyclin binding partners, several of which are upregulated in cancer [36, 37]. AR is known to influence the expression of cell cycle regulatory proteins (e.g., cyclin D1, CDK4, and p21 ^cip1^), but our data suggest that CYP3A5 might have a broader influence on cell cycle regulatory proteins, as several of the regulatory proteins (cyclin B and CHK1) that are not directly regulated by AR are also downregulated after CYP3A5 inhibition.

We observed reduced mRNA expression of cyclin E and CDK2, key regulators of the G1/S transition, in MDAPCa2b cells (AA prostate cancer cell line). This decrease was accompanied by downregulation of the transcription factors E2F1 and E2F2, which control cyclin E expression. Additionally, we found lower mRNA levels of SKP2, a suppressor of p27, leading to increased p27 protein levels, which inhibited the G1/S transition. CDK2 protein levels were downregulated in all three cell lines following treatment with CYP3A5 siRNA and clobetasol. In addition, p27^kip1^ protein (cyclin E2-CDK2 inhibitor) levels were upregulated in all three cell lines treated with siRNA and clobetasol, suggesting that this pathway, involving CYP3A5, may play a central role in blocking G1/S transition [38].

We also found that CYP3A5 siRNA treatment led to downregulation of Cdc25A, a crucial regulator of the G1/S phase transition. The Cdc25 phosphatase family of proteins plays a significant role in cyclin E/A-CDK 2 complex formation as they dephosphorylate Thr14 and Tyr15 residues in CDK2, activating it and allowing it to bind to cyclins. CYP3A5 siRNA-mediated Cdc25A downregulation was compensated for by the downregulation of Chk1 mRNA levels; Chk1 is known to phosphorylate Cdc25A and trigger its ubiquitination and degradation [39]. This interplay between Cdc25A and Chk1 likely contributes to the G1/S transition and affects the cyclin E/CDK2 pair [40]. We observed downregulation of cyclin D1 in all three cell lines treated with CYP3A5 siRNA as well as in MDAPCa2b and LNCaP cells treated with clobetasol, reinforcing our observation of a G1/S block.

Additionally, we investigated the mRNA expression and protein levels of G2/M cyclin-CDK pairs, including CyclinA/CDK1 and CyclinB/CDK1. These genes were downregulated by CYP3A5 siRNA in MDAPCa2b cells. We also observed downregulation of Cyclin B1 protein levels in all three PC cell lines and cyclin A2 protein levels in MDAPCa2b and LNCaP cells. These findings suggest that CYP3A5 siRNA treatment affects the G1/S checkpoint regulation in all cell lines, along with other nodes involved in cell cycle progression.

Our data also shed light on the roles of CHK1 and WEE1 in cell cycle. Although CHK1 and Wee1 can function as tumor suppressors, their overexpression has been observed in many cancers [41, 42]. Our results indicate that CYP3A5 siRNA treatment led to less negative regulation of p-WEE1 at S642, indicating reduced cellular stress. The loss of the G2/M population or the absence of change in the G2/M population can depend on which of the nodes (PLK1, Chk1, Cdc25c, Wee1, and CDK1) predominates.

Another essential component of cell division is DNA replication, which involves the minichromosomal maintenance (MCM) complex, a DNA helicase composed of MCM2-MCM7 proteins. Our findings showed downregulation of the mRNA expression of several MCMs following CYP3A5 siRNA treatment in MDAPCa2b cells. The upregulation of MCM2 to MCM6 is associated with poor prognosis in prostate cancer and liver metastasis [43].

Furthermore, our KEGG analysis revealed differential regulation of genes involved in cellular senescence, DNA replication, and base excision repair, including CALML5, FOXM1, RPA3, HMGB1, FEN1, LIG1, PCNA, BRCA2, and PARPBP [44, 45]. These genes are known to be associated with disease progression in several types of cancers, including prostate cancer [46, 47].

Cell cycle regulation is highly complex, and in many cancer types, it is disrupted, resulting in the aberrant expression of cell cycle proteins. In prostate cancer, CDK4 and CDK6 are often upregulated, leading to hyperactivation of the cyclinD-CDK4/6 axis, overriding Rb-1 negative regulation [48-50]. We used clobetasol propionate, a CYP3A5-specific inhibitor, in combination with palbociclib, an FDA-approved CDK4/6 inhibitor [33, 51]. The combination of a low dose of clobetasol and palbociclib was more effective than either drug alone in killing PC cells, especially MDAPCa2b cells expressing high CYP3A5 protein levels. This synergy is attributed to the distinct targets of the two drugs: palbociclib targets CDK4/6, whereas clobetasol targets the cell cycle regulatory protein CDK2 in all cell lines.

Notably, MDAPCa2b of AA origin expressed high CYP3A5 protein levels (lowest IC_50_ for clobetasol), emphasizing the significance of targeting CYP3A5 in AAs. CYP3A5 promotes AR activation, a key factor in PC growth. Additionally, since CYP3A5 siRNA targets are not dependent on the Rb phosphorylation state, it provides an alternative strategy to target tumors with Rb loss, which do not respond to CDK4/6 inhibitors. CDK2 is frequently upregulated in tumors resistant to CDK4/6 inhibitors, prompting the development of new therapeutic strategies and inhibitors targeting all three CDKs (4, 6, and 2) to overcome resistance. [52-54]. Our combination studies, showing a CI index of less than 0.5 for clobetasol and palbociclib in the AA cell line (MDAPCa2b), support the effectiveness of targeting both CYP3A5/CDK2 and CDK4/6. This strategy may be particularly beneficial for AA men with high CYP3A5 expression, as it also inhibits AR activation.

Our clonogenic assay also demonstrated a significant reduction in cell growth when cells were treated with either clobetasol alone or in combination with palbociclib in all the tested PC lines. The IC_50_ of clobetasol, as determined by the clonogenic assay, was significantly lower than that calculated by the growth assay (methylene blue), particularly in MDAPCa2b cells (800 nM). Because the clonogenic assay measures long-term cell survival, lower clonogenic IC_50_ values suggest a more favorable outcome in a therapeutic setting. This is also particularly important, as it suggests potential therapeutic benefits in castration-resistant prostate cancer (CRPC) cases, including those expressing mutant AR variants such as ARV7 (22RV1).

In summary, our study emphasizes the crucial role of CYP3A5 in regulating prostate cancer cell growth by modulating cell cycle proteins, particularly in African Americans, who often present with aggressive, treatment-resistant diseases. Targeting CYP3A5, especially in combination with CDK4/6 inhibition, shows promise as a therapeutic strategy and can be evaluated with other cell cycle inhibitors. Prednisone, a steroid, is commonly prescribed to patients with prostate cancer, along with abiraterone [55, 56]. An alternative strategy we plan to explore in future studies is to replace prednisone with clobetasol, which can serve as both a steroid and a CYP3A5 inhibitor. Overall, our findings support further investigation of clobetasol propionate as a potential CYP3A5 inhibitor in prostate cancer therapy, particularly in AA patients.

### Conclusion

In this study, we elucidated the effect of CYP3A5 inhibition on prostate cancer cells, leading to a robust cell cycle block at the G1/S checkpoint. RNA sequencing analysis revealed downregulation of multiple critical cell cycle regulatory genes, underscoring the pivotal role of CYP3A5 in modulating cell cycle progression. Furthermore, we observed significant modulation of key cell cycle regulatory proteins, including cyclin D, CDK2, and cyclin B, with consistent emphasis on G1/S checkpoint regulation. CDK2 has emerged as a key molecule affected by both siRNA and clobetasol treatment. Notably, our investigation revealed that the MDAPCa2b cell line, characterized by the highest CYP3A5 expression, displayed the lowest IC_50_ for clobetasol, reinforcing the direct correlation between CYP3A5 levels and treatment response. Notably, the synergistic effects observed with the combination of clobetasol and palbociclib (CDK4/6 inhibitor) were more pronounced in MDAPCa2b cells than in LNCaP cells, highlighting its potential as a therapeutic strategy, especially in African American patients who often exhibit elevated CYP3A5 expression levels. This study underscores the feasibility of targeting CYP3A5 in prostate cancer therapy, with promise for individuals of African American descent expressing high CYP3A5 levels.

## Supporting information

all supplementary figures and tables

## Disclosures

### Ethics statement

N/A (no human subjects/data, tissues, or animals) was used in this study.

### Author contributions

Jeetesh Sharma: Data curation, formal analysis, validation, investigation, visualization, methodology, and writing of the original draft. Imran K. Mohammed: Data curation, formal analysis, validation, investigation, and visualization. Richard L. Tillett: Data curation, formal analysis, methodology, writing– original draft. Jake McLean: Data curation, investigation, and methodology. Shirley Shen: Data curation, investigation, and methodology. Ajay Singh: Resources, investigation, writing–review, and editing. Oscar B. Goodman Jr.: Resources, investigation, writing, reviewing, and editing. Edwin C. Oh: Resources, formal analysis, writing, reviewing, and editing. Ranjana Mitra: Conceptualization, resources, data curation, formal analysis, supervision, funding acquisition, visualization, methodology, writing, project administration, reviewing, and editing.

## Acknowledgement

The authors acknowledge Dr. Aparna Dixit for helping with the review and editing of the manuscript.

