## Supplementary material for "CYP3A5 inhibition causes G1/S blockade and synergizes with CDK4/6 inhibitor to suppress prostate cancer cell growth: Implications in reducing health disparity": all supplementary figures and tables

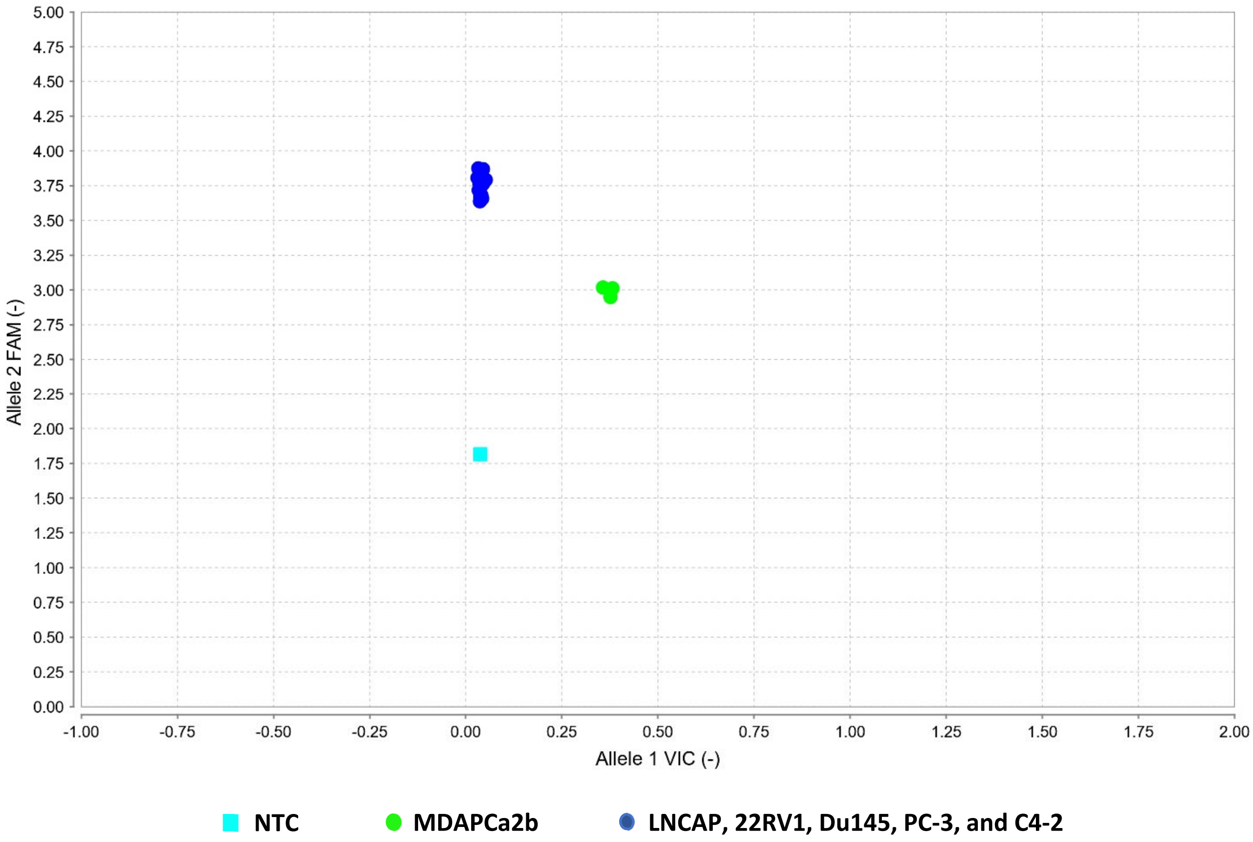


**Figure S1: Genotyping of PC cell lines:** Genomic DNA was isolated from mentioned PC cell lines and 50 ng of genomic DNA was genotyped for detection of wt (*1) and mut (*3) allele of CYP3A5. Allele 1 VIC at x-axis, represent the Wild type (*1) allele and Allele 2 FAM at y-axis, represents mutant (*3) allele. MDAPCa2b have both *1/*3 allele whereas all the other mentioned PCa cell lines were *3/*3 allele. Non-template control (NTC), reaction mixture without DNA template was taken as negative control.

A

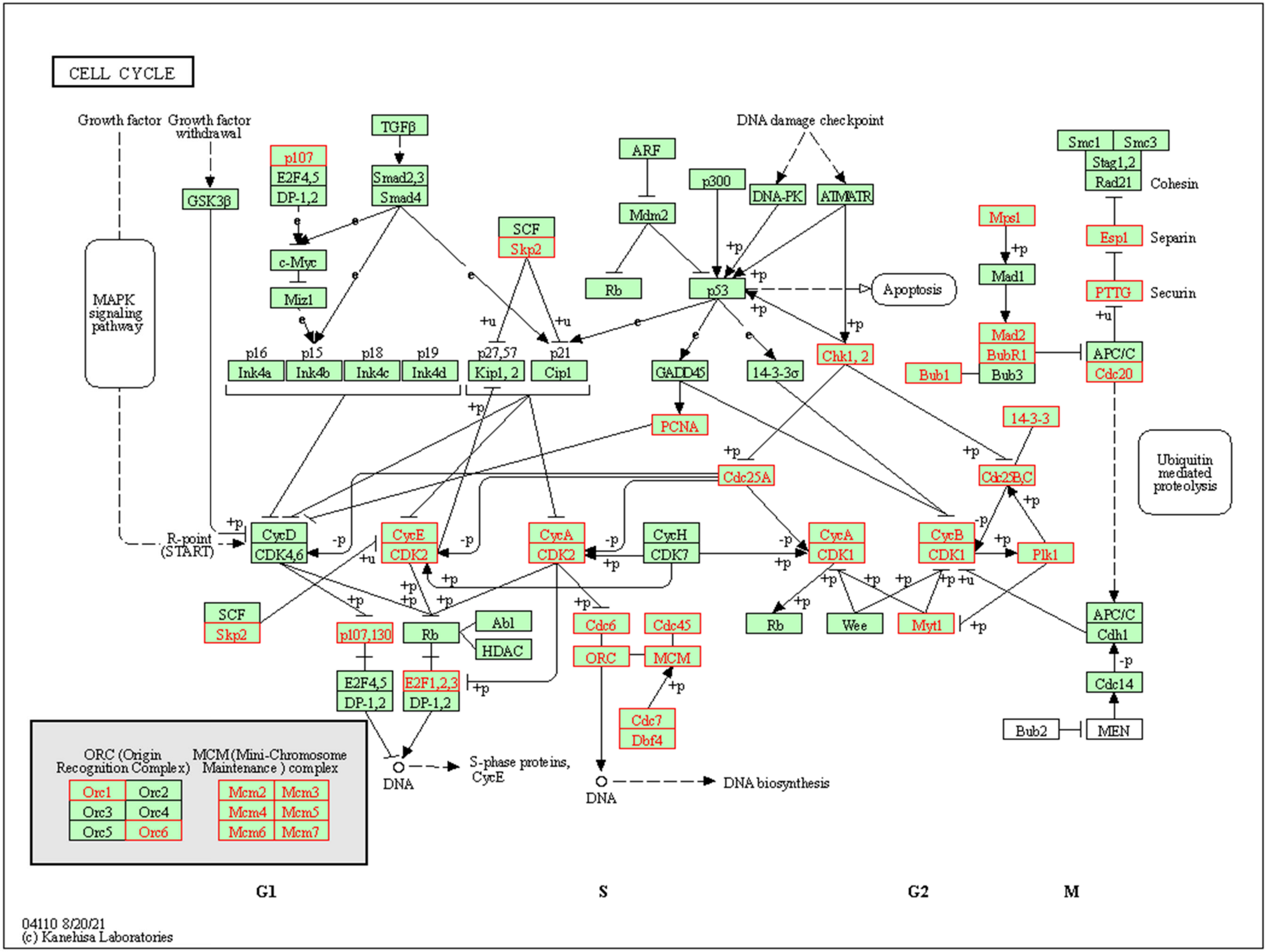
CELL CYCLE
Growth factor
Growth factor withdrawal
MAPK signaling pathway
GSK3β
R-point (START)
p107
E2F4,5
DP-1,2
TGFβ
Smad2,3
Smad4
c-Myc
Miz1
p16
Ink4a
p15
Ink4b
p18
Ink4c
p19
Ink4d
SCF
Skp2
p27,57
Kip1, 2
p21
Cip1
ARF
Mdm2
Rb
p300
DNA damage checkpoint
DNA-PK
ATM/ATR
p53
Apoptosis
GADD45
14-3-3σ
Chk1, 2
PCNA
Cdc25A
CycD
CDK4,6
CycE
CDK2
CycA
CDK2
CycH
CDK7
CycA
CDK1
CycB
CDK1
Plk1
14-3-3
Cdc25B,C
Mps1
Mad1
Mad2
BubR1
Bub3
Bub1
Smc1
Smc3
Stag1,2
Rad21
Cohesin
Esp1
Separin
PTTG
Securin
APC/C
Cdc20
Ubiquitin mediated proteolysis
APC/C
Cdh1
Cdc14
Bub2
MEN
SCF
Skp2
p107,130
E2F4,5
DP-1,2
Rb
Abl
HDAC
E2F1,2,3
DP-1,2
DNA
S-phase proteins, CycE
Cdc6
ORC
Cdc45
MCM
Cdc7
Dbf4
DNA
DNA biosynthesis
Rb
Wee
Myt1
ORC (Origin Recognition Complex)
Orc1
Orc2
Orc3
Orc4
Orc5
Orc6
MCM (Mini-Chromosome Maintenance ) complex
Mcm2
Mcm3
Mcm4
Mcm5
Mcm6
Mcm7
G1
S
G2
M
04110 8/20/21
(c) Kanehisa Laboratories

**B**

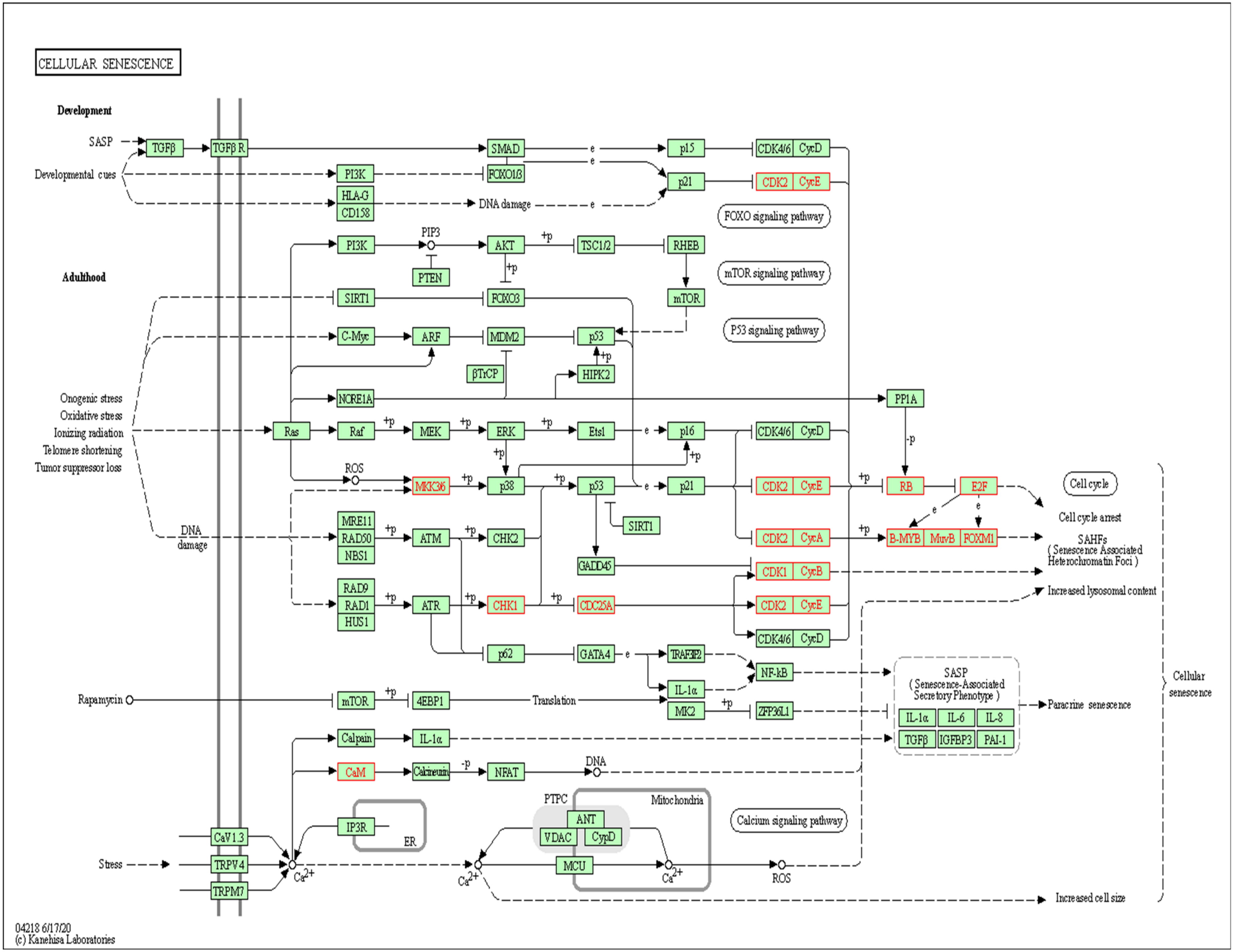
CELLULAR SENESCENCE
Development
SASP
Developmental cues
Adulthood
Onogenic stress
Oxidative stress
Ionizing radiation
Telomere shortening
Tumor suppressor loss
DNA damage
Rapamycin
Stress
TGFβ
TGFβ R
PI3K
HLA-G
CD158
SMAD
FOXO1/3
DNA damage
p15
p21
CDK4/6
CycD
CDK2
CycE
FOXO signaling pathway
mTOR signaling pathway
P53 signaling pathway
PIP3
AKT
PTEN
TSC1/2
RHEB
mTOR
SIRT1
FOXO3
C-Myc
ARF
MDM2
p53
βTrCP
HIPK2
NCRE1A
PP1A
Ras
Raf
MEK
ERK
Ets1
p16
ROS
MKK3/6
p38
RB
E2F
Cell cycle
Cell cycle arrest
MRE11
RAD50
NBS1
ATM
CHK2
CycA
B-MYB
MuvB
FOXM1
SAHFs
( Senescence Associated Heterochromatin Foci )
GADD45
CDK1
CycB
RAD9
RAD1
HUS1
ATR
CHK1
CDC25A
Increased lysosomal content
p62
GATA4
TRAF3IP2
IL-1α
NF-kB
SASP
( Senescence-Associated Secretory Phenotype )
4EBP1
Translation
MK2
ZFP36L1
IL-6
IL-8
Paracrine senescence
Cellular senescence
Calpain
IGFBP3
PAI-1
CaM
Calcineurin
NFAT
DNA
PTPC
Mitochondria
ANT
VDAC
CypD
MCU
IP3R
ER
Calcium signaling pathway
CaV1.3
TRPV4
TRPM7
Ca2+
Increased cell size
04218 6/17/20
(c) Kanehisa Laboratories

C

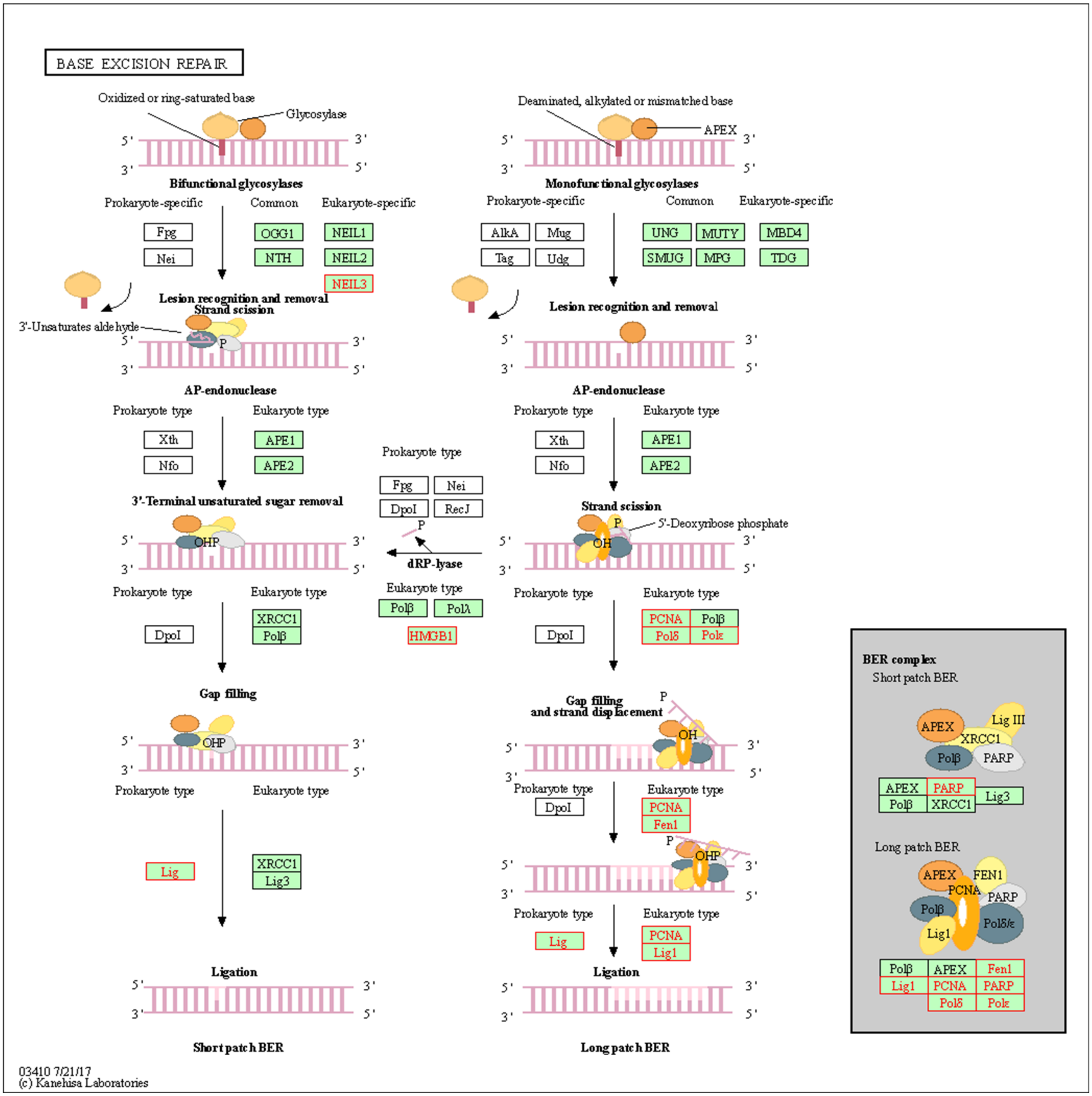

BASE EXCISION REPAIR
Oxidized or ring-saturated base
Glycosylase
Bifunctional glycosylases
Prokaryote-specific
Common
Eukaryote-specific
Fpg
Nei
OGG1
NTH
NEIL1
NEIL2
NEIL3
Lesion recognition and removal
Strand scission
3'-Unsaturates aldehyde
AP-endonuclease
Prokaryote type
Eukaryote type
Xth
Nfo
APE1
APE2
3'-Terminal unsaturated sugar removal
DpoI
XRCC1
Polβ
Gap filling
Lig
Lig3
Ligation
Short patch BER
Deaminated, alkylated or mismatched base
APEX
Monofunctional glycosylases
AlkA
Mug
Tag
Udg
UNG
MUTY
SMUG
MPG
MBD4
TDG
Lesion recognition and removal
Strand scission
5'-Deoxyribose phosphate
Fpg
Nei
DpoI
RecJ
dRP-lyase
Polλ
HMGB1
PCNA
Polδ
Polε
Gap filling
and strand displacement
Fen1
Lig1
Long patch BER
BER complex
Short patch BER
Lig III
PARP
Long patch BER
FEN1
Polδ/ε
03410 7/21/17
(c) Kanehisa Laboratories

**D**

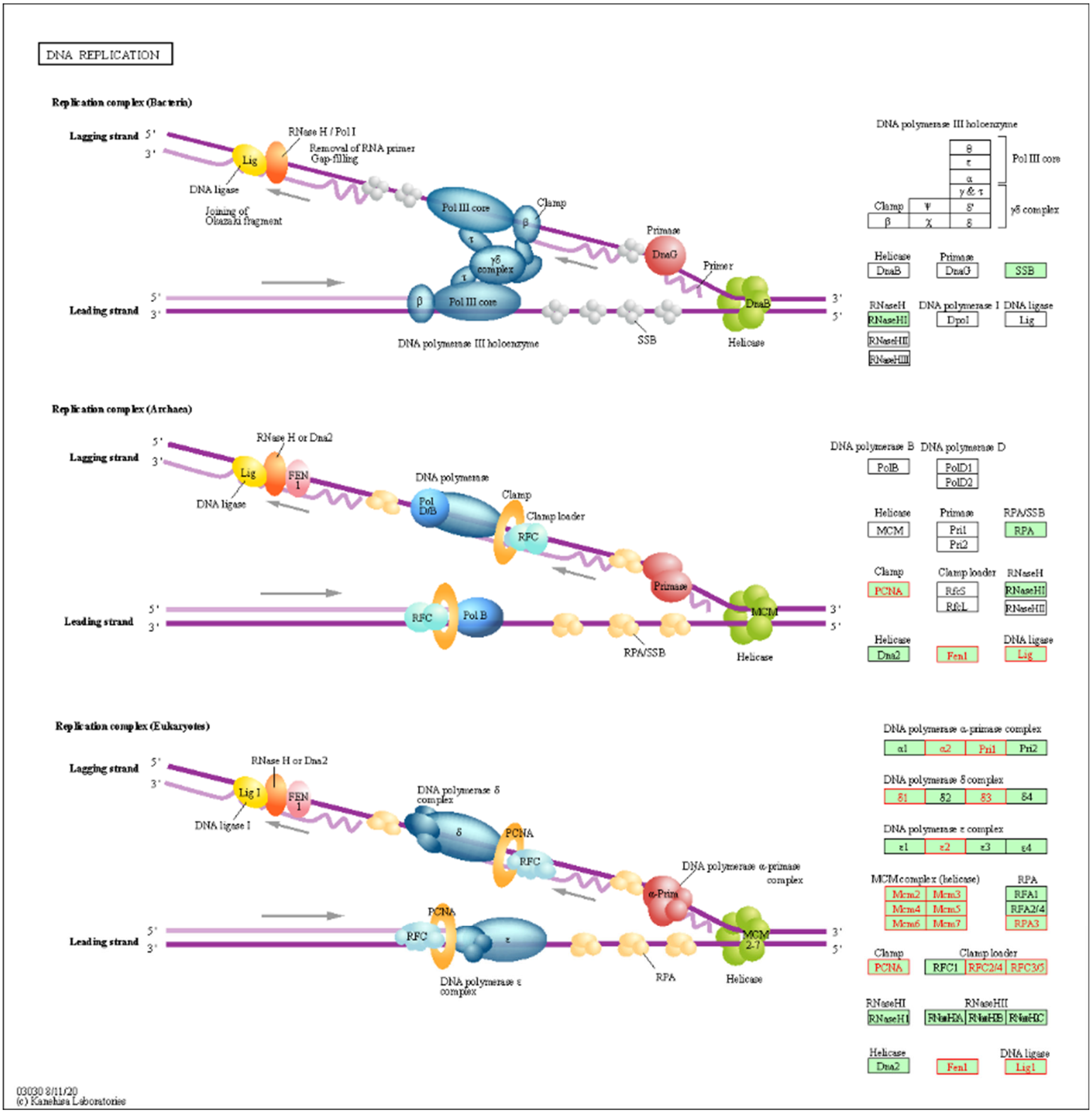

DNA REPLICATION
Replication complex (Bacteria)
Lagging strand
Leading strand
RNase H / Pol I
Removal of RNA primer
Gap-filling
Lig
DNA ligase
Joining of Okazaki fragment
Pol III core
Clamp
γδ complex
Primase
DnaG
Primer
DnaB
DNA polymerase III holoenzyme
SSB
Helicase
DNA polymerase III holoenzyme
Pol III core
γδ complex
Clamp
RNaseH
RNaseHI
RNaseHII
RNaseHIII
DNA polymerase I
DpoI
DNA ligase
Replication complex (Archaea)
RNase H or Dna2
FEN 1
DNA polymerase
Pol D/B
Clamp loader
RFC
Pol B
RPA/SSB
MCM
DNA polymerase B
PolB
DNA polymerase D
PolD1
PolD2
Pri1
Pri2
RPA
PCNA
RfcS
RfcL
Dna2
Fen1
Replication complex (Eukaryotes)
Lig I
DNA ligase I
DNA polymerase δ complex
DNA polymerase α-primase complex
α-Prim
MCM 2-7
DNA polymerase ε complex
MCM complex (helicase)
Mcm2
Mcm3
Mcm4
Mcm5
Mcm6
Mcm7
RFA1
RFA2/4
RPA3
RFC1
RFC2/4
RFC3/5
RNaseH1
RNaseHII
RNaseH2A
RNaseH2B
RNaseH2C
Lig1
03030 8/11/20
(c) Kanehisa Laboratories

E

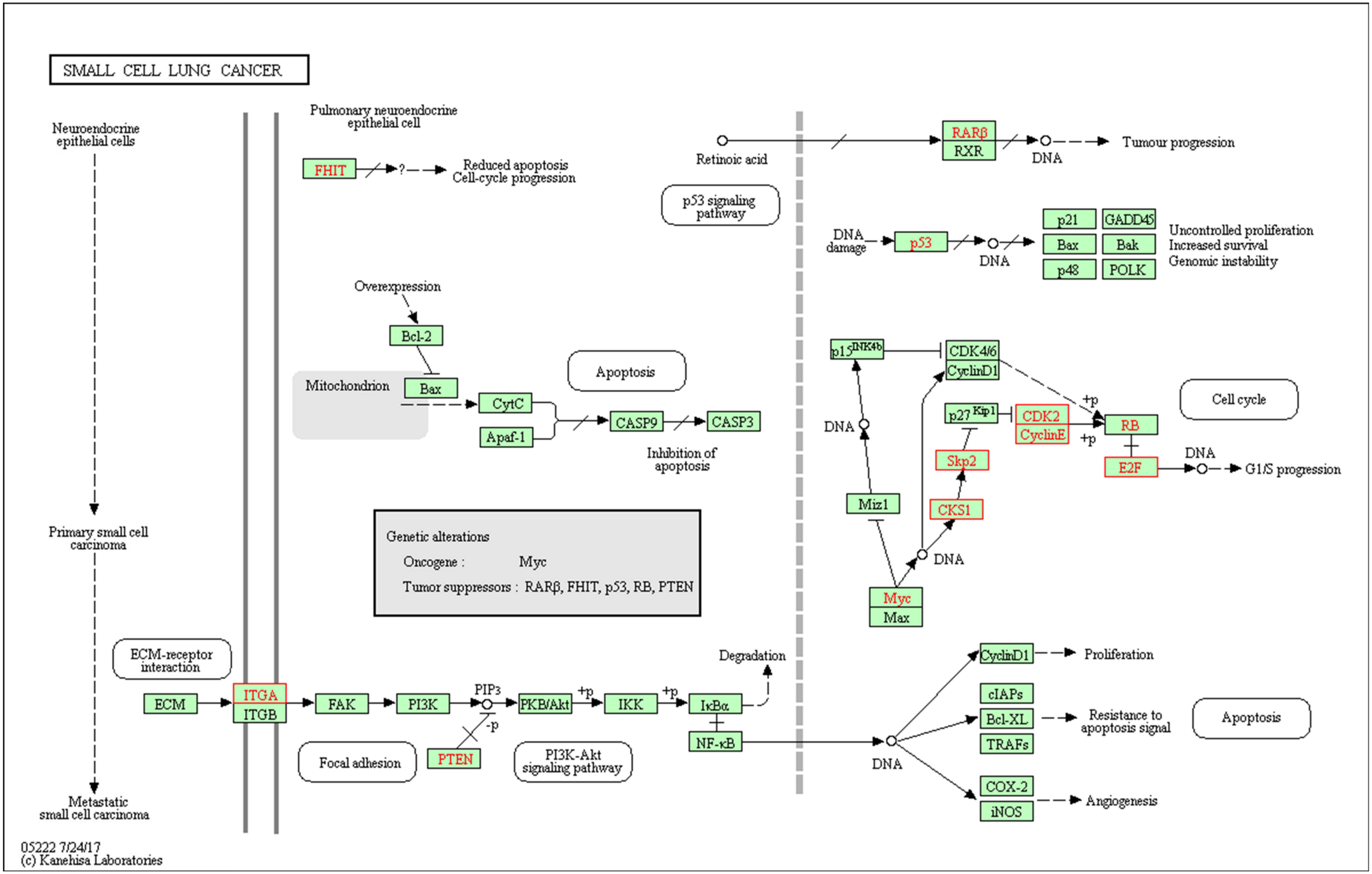
SMALL CELL LUNG CANCER
Neuroendocrine epithelial cells
Primary small cell carcinoma
Metastatic small cell carcinoma
Pulmonary neuroendocrine epithelial cell
FHIT
?
Reduced apoptosis
Cell-cycle progression
Retinoic acid
RARβ
RXR
DNA
Tumour progression
p53 signaling pathway
DNA damage
p53
p21
GADD45
Bax
Bak
p48
POLK
Uncontrolled proliferation
Increased survival
Genomic instability
Overexpression
Bcl-2
Mitochondrion
Bax
CytC
Apaf-1
Apoptosis
CASP9
CASP3
Inhibition of apoptosis
p15INK4b
CDK4/6
CyclinD1
p27Kip1
CDK2
CyclinE
+p
RB
E2F
G1/S progression
Cell cycle
Skp2
CKS1
Miz1
Myc
Max
Genetic alterations
Oncogene : Myc
Tumor suppressors : RARβ, FHIT, p53, RB, PTEN
ECM-receptor interaction
ECM
ITGA
ITGB
FAK
PI3K
PIP3
-p
PTEN
PKB/Akt
IKK
IκBα
NF-κB
Degradation
Focal adhesion
PI3K-Akt signaling pathway
Proliferation
cIAPs
Bcl-XL
TRAFs
Resistance to apoptosis signal
COX-2
iNOS
Angiogenesis
05222 7/24/17
(c) Kanehisa Laboratories

F

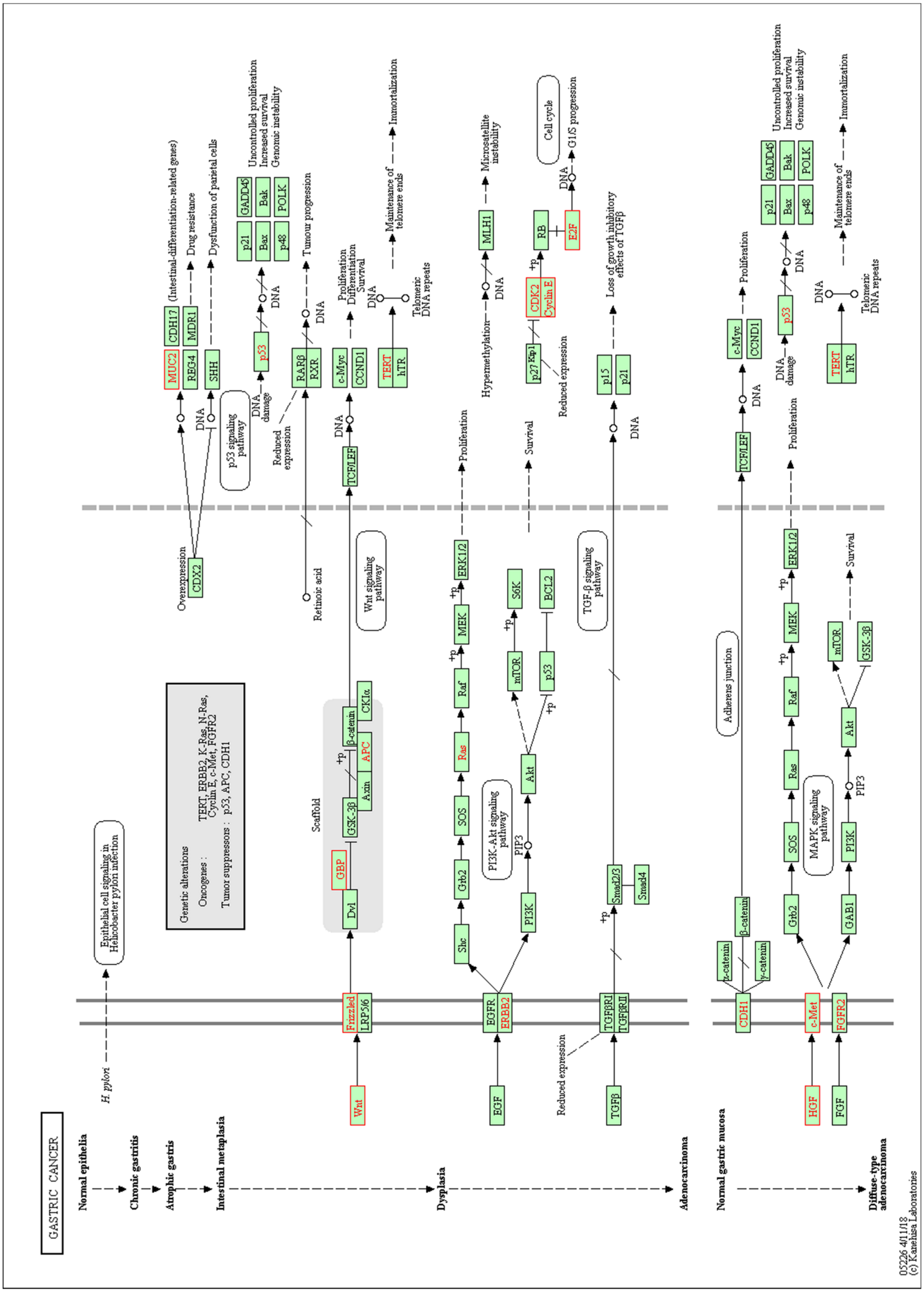


**Figure S2:** KEGG pathway analysis suggested the involvement of four significant pathways (Cell cycle (A), DNA replication (B), Base excision repair (C), Cellular senescence (D), small cell lung cancer (E), and Gastric cancer (F)) that are relevant to our study with *p* value less than 0.01. A list of the gene involved in each pathway is enlisted in supplementary table 1.

**Table S1.** Gene list with the enrichment score for the significant pathway after GSEA.

| Gene | fold change | Adj *p*-value | Gene | fold change | Adj *p*-value |
|---|---|---|---|---|---|
| **Oocyte meiosis** | | | **Fanconi anemia pathway** | | |
| FBXO43 | -5.257602767 | 0.032406886 | RMI2 | -3.89720874 | 1.34E-21 |
| CDK2 | -4.309184682 | 1.43E-21 | RAD51 | -3.231969271 | 1.69E-16 |
| AURKA | -4.249556636 | 2.92E-31 | FANCA | -2.85006138 | 3.92E-18 |
| PTTG1 | -3.6423789 | 8.90E-08 | BLM | -2.778046773 | 5.19E-08 |
| CCNB2 | -3.464863241 | 3.96E-22 | EME1 | -2.761896002 | 1.27E-09 |
| CCNB1 | -3.446825772 | 4.94E-30 | FANCI | -2.757523098 | 5.54E-18 |
| PLK1 | -3.421834419 | 2.28E-36 | FANCD2 | -2.739833704 | 6.78E-11 |
| CDC20 | -3.279812089 | 6.05E-14 | UBE2T | -2.709242269 | 2.43E-11 |
| CDK1 | -3.235539039 | 3.55E-12 | FANCG | -2.61279915 | 3.93E-17 |
| PKMYT1 | -3.216121264 | 1.01E-12 | BRCA2 | -2.6097387 | 0.003682393 |
| **Cellular senescence** | | | **Human immunodeficiency virus 1 infection** | | |
| CDK2 | -4.309184682 | 1.43E-21 | CHEK1 | -3.530906632 | 3.27E-11 |
| CCNE1 | -2.697400472 | 3.22E-10 | CCNB2 | -3.464863241 | 3.96E-22 |
| CCNA2 | -3.329403972 | 2.17E-09 | CCNB1 | -3.446825772 | 4.94E-30 |
| CDK1 | -3.235539039 | 3.55E-12 | PAK1 | -3.289690467 | 9.11E-29 |
| CCNB2 | -3.464863241 | 3.96E-22 | CDK1 | -3.235539039 | 3.55E-12 |
| CCNB1 | -3.446825772 | 4.94E-30 | CALML5 | -3.02681161 | 0.018292764 |
| CHEK1 | -3.530906632 | 3.27E-11 | APOBEC3B | -2.97650815 | 8.08E-13 |
| CDC25A | -2.515801445 | 9.04E-13 | CDC25C | -2.921396697 | 2.07E-10 |
| MYBL2 | -3.795651257 | 4.29E-24 | | | |
| LIN9 | -3.314934451 | 0.0000117 | **Gastric cancer** | | |
| FOXM1 | -3.903361987 | 1E-33 | CDK2 | -4.309184682 | 1.43E-21 |
| CALML5 | -3.02681161 | 0.018293 | MET | -4.126482197 | 2.84E-06 |
| | | | MUC2 | -3.628022339 | 1.63E-18 |
| **Progesterone-mediated oocyte maturation** | | | WNT11 | -3.090298014 | 0.0422169 |
| CDK2 | -4.309184682 | 1.43E-21 | WNT10B | -2.970218471 | 1.58E-04 |
| AURKA | -4.249556636 | 2.92E-31 | CCNE1 | -2.697400472 | 3.22E-10 |
| CCNB2 | -3.464863241 | 3.96E-22 | E2F1 | -2.62421406 | 2.31E-09 |
| CCNB1 | -3.446825772 | 4.94E-30 | E2F2 | -2.498381404 | 9.14E-12 |
| PLK1 | -3.421834419 | 2.28E-36 | WNT5A | -2.304307917 | 0.0186456 |
| CCNA2 | -3.329403972 | 2.17E-09 | FRAT2 | -2.250605535 | 2.72E-07 |
| CDK1 | -3.235539039 | 3.55E-12 | | | |
| PKMYT1 | -3.216121264 | 1.01E-12 | **Base excision repair** | | |
| BUB1 | -2.991891852 | 8.88E-08 | PARPBP | -3.35406 | 0.0000242 |
| CDC25C | -2.921396697 | 2.07E-10 | FEN1 | -3.04291 | 2.61E-11 |
| | | | PCNA | -2.21665 | 1.93E-10 |
| **DNA replication** | | | LIG1 | -2.34271 | 9.16E-18 |
| FEN1 | -3.042910933 | 2.61E-11 | HMGB1 | -2.11472 | 8.7E-12 |
| PCNA | -2.216648062 | 1.93E-10 | NEIL3 | -2.84346 | 0.00000714 |
| LIG1 | -2.342712108 | 9.16E-18 | BRCA2 | -2.60974 | 0.0036824 |
| MCM5 | -3.291755743 | 3.16E-24 | | | |
| MCM2 | -2.646965872 | 2.2E-15 | **Small cell lung cancer** | | |
| MCM6 | -2.081600169 | 2.89E-12 | CDK2 | -4.309184682 | 1.43E-21 |
| MCM4 | -2.036839662 | 0.00000879 | CKS2 | -3.203651129 | 3.21E-19 |
| POLE2 | -2.786136946 | 5.76E-13 | CKS1B | -2.999359093 | 2.83E-08 |

| | | | | | |
|---|---|---|---|---|---|
| POLA2 | -2.262296687 | 0.049998 | CCNE1 | -2.697400472 | 3.22E-10 |
| POLD1 | -2.216426223 | 1.02E-11 | E2F1 | -2.62421406 | 2.31E-09 |
| POLD3 | -1.925030136 | 0.00000195 | SKP2 | -2.598134414 | 1.94E-07 |
| PRIM1 | -1.997709087 | 0.00000446 | E2F2 | -2.498381404 | 9.14E-12 |
| MCM7 | -1.841836264 | 0.00001 | ITGA6 | -2.312049246 | 0.015501732 |
| MCM3 | -1.974952859 | 6.38E-14 | | | |
| RPA3 | -1.882443986 | 0.001398302 | **Steroid hormone biosynthesis** | | |
| RPP25 | -2.179774259 | 0.002825758 | AKR1D1 | 8.953356908 | 0.04186974 |
| | | | CYP1B1 | 8.339150505 | 2.00E-12 |
| **Human T-cell leukemia virus 1 infection** | | | CYP11A1 | 4.813114458 | 0.012256102 |
| MYBL2 | -3.795651257 | 4.29E-24 | UGT2B11 | 4.387159812 | 4.53E-17 |
| PTTG1 | -3.6423789 | 8.90E-08 | UGT2B4 | 3.246455716 | 9.71E-08 |
| CHEK1 | -3.530906632 | 3.27E-11 | UGT2B10 | 2.91595143 | 3.33E-04 |
| CCNB2 | -3.464863241 | 3.96E-22 | | | |
| CDC20 | -3.279812089 | 6.05E-14 | **Retinol metabolism** | | |
| WNT11 | -3.090298014 | 0.0422169 | UGT2B11 | 4.387159812 | 4.53E-17 |
| WNT10B | -2.970218471 | 1.58E-04 | RDH12 | 4.35618736 | 0.02693986 |
| MAD2L1 | -2.900871214 | 4.31E-07 | UGT2B4 | 3.246455716 | 9.71E-08 |
| BUB1B | -2.841828213 | 5.03E-05 | RDH16 | 2.938267686 | 6.56E-06 |
| POLE2 | -2.786136946 | 5.76E-13 | UGT2B10 | 2.91595143 | 3.33E-04 |
| | | | ALDH1A1 | 2.323591519 | 0.022669571 |
| | | | UGT2B15 | 2.091049609 | 1.87E-09 |

**Table S2:** List of 1814 differentially expressed genes after CYP3A5 siRNA treatment in MDAPCa2b PCa cell.

| Gene | GeneID | log2FoldChange | pvalue | padj | chromosome |
|---|---|---|---|---|---|
| IGFBP4 | ENSG00000141753 | -5.055869819 | 1.04E-04 | 0.002066068 | 17 |
| LINC01482 | ENSG00000267659 | -4.585143492 | 6.88E-04 | 0.010131088 | 17 |
| GPX8 | ENSG00000164294 | -3.857304404 | 5.33E-07 | 2.14E-05 | 5 |
| TFCP2L1 | ENSG00000115112 | -3.734456234 | 0.001984069 | 0.023638759 | 2 |
| ACTL8 | ENSG00000117148 | -3.549350666 | 3.85E-04 | 0.006220649 | 1 |
| FAM20A | ENSG00000108950 | -3.435008012 | 7.82E-04 | 0.011200142 | 17 |
| IL37 | ENSG00000125571 | -3.114529614 | 1.63E-21 | 5.74E-19 | 2 |
| GLO1 | ENSG00000124767 | -3.112894583 | 5.22E-76 | 8.99E-72 | 6 |
| NES | ENSG00000132688 | -3.087123521 | 5.95E-07 | 2.32E-05 | 1 |
| FCGRT | ENSG00000104870 | -2.944591058 | 0.001308669 | 0.016899802 | 19 |
| DHRS2 | ENSG00000100867 | -2.860335176 | 6.19E-10 | 4.37E-08 | 14 |
| | ENSG00000254854 | -2.818860661 | 1.94E-07 | 8.50E-06 | 11 |
| RFTN1 | ENSG00000131378 | -2.793624096 | 1.82E-48 | 1.57E-44 | 3 |
| MGP | ENSG00000111341 | -2.695303983 | 1.03E-08 | 5.82E-07 | 12 |
| LINC01962 | ENSG00000248473 | -2.638696093 | 0.003202975 | 0.03434225 | 5 |
| ETNK2 | ENSG00000143845 | -2.60912897 | 8.95E-07 | 3.28E-05 | 1 |
| | ENSG00000284738 | -2.542467761 | 0.003280734 | 0.034882364 | 1 |
| EMID1 | ENSG00000186998 | -2.511954557 | 2.14E-06 | 7.06E-05 | 22 |
| FBXO43 | ENSG00000156509 | -2.394405145 | 0.002974491 | 0.032406886 | 8 |
| USP18 | ENSG00000184979 | -2.357093271 | 5.96E-25 | 3.02E-22 | 22 |
| MIR1244-2 | ENSG00000283498 | -2.325848189 | 1.86E-10 | 1.46E-08 | 5 |

| | | | | | |
|---|---|---|---|---|---|
| | ENSG00000286729 | -2.321701175 | 0.00483645 | 0.047089739 | 3 |
| RERG | ENSG00000134533 | -2.205565591 | 1.32E-10 | 1.05E-08 | 12 |
| TEAD2 | ENSG00000074219 | -2.187658945 | 1.30E-29 | 1.49E-26 | 19 |
| GAS2L3 | ENSG00000139354 | -2.187080022 | 6.15E-06 | 1.81E-04 | 12 |
| INSIG1 | ENSG00000186480 | -2.171422738 | 7.60E-24 | 3.19E-21 | 7 |
| APOC1 | ENSG00000130208 | -2.164993308 | 7.98E-06 | 2.29E-04 | 19 |
| EBF2 | ENSG00000221818 | -2.138615855 | 0.00357943 | 0.037207916 | 8 |
| TK1 | ENSG00000167900 | -2.137094448 | 6.45E-17 | 1.42E-14 | 17 |
| | ENSG00000275185 | -2.131073032 | 0.00238535 | 0.027301477 | 17 |
| LDAH | ENSG00000118961 | -2.126883302 | 2.23E-04 | 0.003936014 | 2 |
| PIMREG | ENSG00000129195 | -2.12429754 | 2.54E-25 | 1.36E-22 | 17 |
| APLN | ENSG00000171388 | -2.111250852 | 3.17E-09 | 1.97E-07 | X |
| C5orf34 | ENSG00000172244 | -2.10825864 | 1.09E-10 | 8.83E-09 | 5 |
| CDK2 | ENSG00000123374 | -2.107414931 | 3.23E-24 | 1.43E-21 | 12 |
| TROAP | ENSG00000135451 | -2.102890501 | 1.26E-28 | 1.20E-25 | 12 |
| AURKA | ENSG00000087586 | -2.08731233 | 1.36E-34 | 2.92E-31 | 20 |
| NMU | ENSG00000109255 | -2.08381107 | 6.89E-07 | 2.62E-05 | 4 |
| ANGPTL2 | ENSG00000136859 | -2.079839979 | 2.00E-11 | 1.83E-09 | 9 |
| | ENSG00000259020 | -2.072175632 | 0.003075397 | 0.033295523 | 14 |
| CENPE | ENSG00000138778 | -2.069486358 | 2.60E-08 | 1.36E-06 | 4 |
| RUNX1T1 | ENSG00000079102 | -2.062786371 | 0.00266312 | 0.029691024 | 8 |
| TNS4 | ENSG00000131746 | -2.061519857 | 3.50E-12 | 3.72E-10 | 17 |
| A2M | ENSG00000175899 | -2.060550875 | 4.49E-10 | 3.27E-08 | 12 |
| SHISA2 | ENSG00000180730 | -2.048378171 | 6.20E-07 | 2.41E-05 | 13 |
| OIP5 | ENSG00000104147 | -2.045050849 | 9.47E-21 | 2.91E-18 | 15 |
| MET | ENSG00000105976 | -2.044912416 | 5.88E-08 | 2.84E-06 | 7 |
| CYP3A5 | ENSG00000106258 | -2.030145105 | 3.39E-12 | 3.62E-10 | 7 |
| LGALS1 | ENSG00000100097 | -2.029214719 | 4.18E-05 | 9.56E-04 | 22 |
| WIPF3 | ENSG00000122574 | -2.027861834 | 2.80E-04 | 0.004770267 | 7 |
| SP4 | ENSG00000105866 | -2.023605119 | 4.69E-04 | 0.007365968 | 7 |
| CENPK | ENSG00000123219 | -2.008824062 | 2.12E-10 | 1.62E-08 | 5 |
| CAVIN1 | ENSG00000177469 | -2.000116444 | 4.35E-11 | 3.76E-09 | 17 |
| CDKN3 | ENSG00000100526 | -1.984306927 | 2.30E-27 | 1.98E-24 | 14 |
| FOXM1 | ENSG00000111206 | -1.96471726 | 3.50E-37 | 1.00E-33 | 12 |
| RMI2 | ENSG00000175643 | -1.962441206 | 2.95E-24 | 1.34E-21 | 16 |
| TMEM109 | ENSG00000110108 | -1.961903354 | 5.11E-30 | 6.28E-27 | 11 |
| BIRC5 | ENSG00000089685 | -1.954785666 | 5.64E-21 | 1.80E-18 | 17 |
| PRC1 | ENSG00000198901 | -1.947110971 | 5.92E-11 | 5.02E-09 | 15 |
| RDM1 | ENSG00000278023 | -1.937878516 | 7.48E-09 | 4.39E-07 | 17 |
| POLQ | ENSG00000051341 | -1.935886571 | 4.75E-09 | 2.86E-07 | 3 |
| MIR1244-3 | ENSG00000283429 | -1.925558795 | 1.21E-06 | 4.24E-05 | 12 |
| MYBL2 | ENSG00000101057 | -1.924347444 | 5.49E-27 | 4.29E-24 | 20 |
| STMN1 | ENSG00000117632 | -1.924091769 | 4.59E-39 | 1.58E-35 | 1 |
| SKA3 | ENSG00000165480 | -1.919055663 | 2.03E-18 | 5.00E-16 | 13 |
| PEPD | ENSG00000124299 | -1.915682609 | 6.74E-26 | 4.00E-23 | 19 |
| FAM83D | ENSG00000101447 | -1.907439639 | 9.18E-10 | 6.30E-08 | 20 |
| KLK1 | ENSG00000167748 | -1.906565823 | 1.62E-10 | 1.27E-08 | 19 |
| CENPW | ENSG00000203760 | -1.902807753 | 1.69E-08 | 9.23E-07 | 6 |
| PRR11 | ENSG00000068489 | -1.899026186 | 5.22E-19 | 1.34E-16 | 17 |
| GTSE1 | ENSG00000075218 | -1.898589278 | 5.59E-21 | 1.80E-18 | 22 |
| NUF2 | ENSG00000143228 | -1.87936296 | 2.43E-21 | 8.38E-19 | 1 |
| TACR3 | ENSG00000169836 | -1.873728478 | 3.44E-06 | 1.08E-04 | 4 |
| | ENSG00000267698 | -1.868975937 | 1.83E-04 | 0.003349287 | 19 |
| PTTG1 | ENSG00000164611 | -1.864881007 | 1.33E-09 | 8.90E-08 | 5 |
| ASPM | ENSG00000066279 | -1.861178183 | 5.79E-08 | 2.81E-06 | 1 |
| MUC2 | ENSG00000198788 | -1.859183339 | 4.93E-21 | 1.63E-18 | 11 |
| TNFAIP8L1 | ENSG00000185361 | -1.847709504 | 7.66E-14 | 1.03E-11 | 19 |
| TRIM7 | ENSG00000146054 | -1.844808718 | 2.88E-09 | 1.81E-07 | 5 |

| | | | | | |
|---|---|---|---|---|---|
| | ENSG00000287566 | -1.840446252 | 2.29E-12 | 2.51E-10 | 10 |
| NKAP | ENSG00000101882 | -1.834151886 | 1.77E-26 | 1.13E-23 | X |
| CHEK1 | ENSG00000149554 | -1.820038672 | 2.61E-13 | 3.27E-11 | 11 |
| CEP55 | ENSG00000138180 | -1.811521351 | 6.18E-09 | 3.69E-07 | 10 |
| DEPDC1 | ENSG00000024526 | -1.810681284 | 3.89E-07 | 1.60E-05 | 1 |
| FBLN7 | ENSG00000144152 | -1.81048364 | 0.003790316 | 0.038976405 | 2 |
| LBR | ENSG00000143815 | -1.806106103 | 8.64E-07 | 3.19E-05 | 1 |
| LINC01836 | ENSG00000267530 | -1.802241251 | 7.75E-05 | 0.001603107 | 19 |
| ACBD7 | ENSG00000176244 | -1.799982803 | 6.35E-04 | 0.009528053 | 10 |
| CCNB2 | ENSG00000157456 | -1.79279841 | 8.06E-25 | 3.96E-22 | 15 |
| HJURP | ENSG00000123485 | -1.79017225 | 1.54E-26 | 1.02E-23 | 2 |
| MIR924HG | ENSG00000267374 | -1.789581198 | 7.50E-10 | 5.21E-08 | 18 |
| CCNB1 | ENSG00000134057 | -1.785268376 | 2.87E-33 | 4.94E-30 | 5 |
| | ENSG00000268034 | -1.783661507 | 0.002872815 | 0.031498495 | 19 |
| SLC25A10 | ENSG00000183048 | -1.778386285 | 5.47E-07 | 2.16E-05 | 17 |
| PLK1 | ENSG00000166851 | -1.77476995 | 5.30E-40 | 2.28E-36 | 16 |
| PBK | ENSG00000168078 | -1.773747668 | 3.92E-19 | 1.02E-16 | 8 |
| | ENSG00000276672 | -1.771767288 | 0.001088144 | 0.014554239 | 13 |
| STXBP6 | ENSG00000168952 | -1.770554618 | 3.09E-04 | 0.005163809 | 14 |
| AURKB | ENSG00000178999 | -1.769436188 | 1.18E-12 | 1.39E-10 | 17 |
| MTFR2 | ENSG00000146410 | -1.768874648 | 1.01E-16 | 2.12E-14 | 6 |
| GPSM2 | ENSG00000121957 | -1.761264968 | 8.18E-17 | 1.76E-14 | 1 |
| | ENSG00000225173 | -1.755405238 | 0.001057699 | 0.014219094 | 6 |
| TENT5A | ENSG00000112773 | -1.753876708 | 1.19E-05 | 3.25E-04 | 6 |
| HMGB2 | ENSG00000164104 | -1.752911501 | 1.35E-44 | 7.74E-41 | 4 |
| RHBDL3 | ENSG00000141314 | -1.752366368 | 2.79E-04 | 0.004766049 | 17 |
| UBE2C | ENSG00000175063 | -1.7516867 | 1.31E-16 | 2.68E-14 | 20 |
| PARPBP | ENSG00000185480 | -1.745909109 | 6.23E-07 | 2.42E-05 | 12 |
| TFF1 | ENSG00000160182 | -1.74334527 | 8.64E-08 | 4.05E-06 | 21 |
| PCSK9 | ENSG00000169174 | -1.741652336 | 7.74E-15 | 1.23E-12 | 1 |
| CENPA | ENSG00000115163 | -1.739516747 | 1.74E-19 | 4.60E-17 | 2 |
| DZIP1L | ENSG00000158163 | -1.738093751 | 0.001093818 | 0.014618781 | 3 |
| H2AC20 | ENSG00000184260 | -1.736976767 | 3.81E-04 | 0.00616962 | 1 |
| | ENSG00000268119 | -1.73656066 | 2.22E-04 | 0.003935704 | 19 |
| CCNA2 | ENSG00000145386 | -1.73526393 | 2.42E-11 | 2.17E-09 | 4 |
| ORAI1 | ENSG00000276045 | -1.73460897 | 1.74E-09 | 1.13E-07 | 12 |
| GPSM3 | ENSG00000213654 | -1.733150397 | 1.68E-05 | 4.37E-04 | 6 |
| LIN9 | ENSG00000183814 | -1.728980343 | 2.77E-07 | 1.17E-05 | 1 |
| TOX2 | ENSG00000124191 | -1.728068352 | 1.20E-04 | 0.002359475 | 20 |
| FAM72B | ENSG00000188610 | -1.723419526 | 2.66E-10 | 2.01E-08 | 1 |
| CDC6 | ENSG00000094804 | -1.720819 | 1.33E-26 | 9.13E-24 | 17 |
| | ENSG00000187951 | -1.720288987 | 1.16E-04 | 0.002275891 | 15 |
| PIF1 | ENSG00000140451 | -1.719003381 | 5.50E-10 | 3.92E-08 | 15 |
| MCM5 | ENSG00000100297 | -1.718857288 | 3.86E-27 | 3.16E-24 | 22 |
| PAK1 | ENSG00000149269 | -1.717951845 | 5.82E-32 | 9.11E-29 | 11 |
| E2F8 | ENSG00000129173 | -1.716977254 | 6.86E-04 | 0.010116221 | 11 |
| CDC20 | ENSG00000117399 | -1.713613161 | 3.06E-16 | 6.05E-14 | 1 |
| IFI27L2 | ENSG00000119632 | -1.706088073 | 5.12E-06 | 1.54E-04 | 14 |
| NCAPH | ENSG00000121152 | -1.702789924 | 2.65E-15 | 4.61E-13 | 2 |
| GALNTL6 | ENSG00000174473 | -1.700428002 | 8.38E-07 | 3.11E-05 | 4 |
| DIAPH3 | ENSG00000139734 | -1.694349505 | 2.99E-08 | 1.55E-06 | 13 |
| CDK1 | ENSG00000170312 | -1.694006084 | 2.39E-14 | 3.55E-12 | 10 |
| RAD51 | ENSG00000051180 | -1.692413481 | 6.68E-19 | 1.69E-16 | 15 |
| PHF19 | ENSG00000119403 | -1.689860323 | 8.68E-23 | 3.40E-20 | 9 |
| PRKG1 | ENSG00000185532 | -1.688612042 | 1.95E-07 | 8.54E-06 | 10 |
| ASZ1 | ENSG00000154438 | -1.687108033 | 2.94E-05 | 7.07E-04 | 7 |
| PKMYT1 | ENSG00000127564 | -1.685321804 | 6.22E-15 | 1.01E-12 | 16 |
| ZWINT | ENSG00000122952 | -1.682380652 | 3.45E-30 | 4.56E-27 | 10 |

| | | | | | |
|---|---|---|---|---|---|
| NRM | ENSG00000137404 | -1.680891136 | 2.04E-10 | 1.57E-08 | 6 |
| CKS2 | ENSG00000123975 | -1.67971705 | 8.94E-22 | 3.21E-19 | 9 |
| SNHG26 | ENSG00000228649 | -1.679345578 | 0.001635859 | 0.020346585 | 7 |
| ARHGAP11A | ENSG00000198826 | -1.675945566 | 7.19E-05 | 0.001508343 | 15 |
| UCP2 | ENSG00000175567 | -1.675756954 | 2.13E-09 | 1.36E-07 | 11 |
| CERCAM | ENSG00000167123 | -1.67309493 | 1.04E-05 | 2.87E-04 | 9 |
| CKAP2L | ENSG00000169607 | -1.672858502 | 5.51E-08 | 2.70E-06 | 2 |
| MELTF | ENSG00000163975 | -1.669850798 | 2.05E-12 | 2.27E-10 | 3 |
| CENPU | ENSG00000151725 | -1.664156762 | 7.12E-27 | 5.33E-24 | 4 |
| RECQL4 | ENSG00000160957 | -1.657769955 | 8.97E-12 | 8.73E-10 | 8 |
| NUDT1 | ENSG00000106268 | -1.657152293 | 2.95E-05 | 7.08E-04 | 7 |
| PLK4 | ENSG00000142731 | -1.655681175 | 4.43E-06 | 1.36E-04 | 4 |
| DLGAP5 | ENSG00000126787 | -1.648464952 | 9.00E-12 | 8.73E-10 | 14 |
| FLRT3 | ENSG00000125848 | -1.642342767 | 5.20E-04 | 0.008069353 | 20 |
| CDT1 | ENSG00000167513 | -1.637150236 | 6.78E-10 | 4.76E-08 | 16 |
| NEK2 | ENSG00000117650 | -1.635791262 | 8.59E-15 | 1.36E-12 | 1 |
| KIFC1 | ENSG00000237649 | -1.630092133 | 9.80E-27 | 7.03E-24 | 6 |
| CCDC150 | ENSG00000144395 | -1.629497884 | 3.02E-06 | 9.61E-05 | 2 |
| WNT11 | ENSG00000085741 | -1.627745972 | 0.004190661 | 0.0422169 | 11 |
| APOE | ENSG00000130203 | -1.621578558 | 4.17E-05 | 9.56E-04 | 19 |
| SASS6 | ENSG00000156876 | -1.61766324 | 4.59E-04 | 0.007220687 | 1 |
| TACC3 | ENSG00000013810 | -1.617480466 | 5.85E-12 | 5.96E-10 | 4 |
| SLC30A3 | ENSG00000115194 | -1.615570636 | 6.16E-05 | 0.001317181 | 2 |
| KIF18A | ENSG00000121621 | -1.611771546 | 1.70E-07 | 7.49E-06 | 11 |
| PAQR3 | ENSG00000163291 | -1.610039426 | 0.003579313 | 0.037207916 | 4 |
| HMMR | ENSG00000072571 | -1.608965461 | 2.04E-06 | 6.79E-05 | 5 |
| FEN1 | ENSG00000168496 | -1.605452106 | 2.03E-13 | 2.61E-11 | 11 |
| TOMM22 | ENSG00000100216 | -1.602745635 | 8.15E-10 | 5.63E-08 | 22 |
| TSPAN8 | ENSG00000127324 | -1.602031958 | 2.11E-15 | 3.75E-13 | 12 |
| CALML5 | ENSG00000178372 | -1.597798884 | 0.001435664 | 0.018292764 | 10 |
| FUOM | ENSG00000148803 | -1.593769298 | 3.85E-04 | 0.006220649 | 10 |
| KIF2C | ENSG00000142945 | -1.592754593 | 3.37E-25 | 1.76E-22 | 1 |
| PEBP4 | ENSG00000134020 | -1.589618184 | 5.12E-07 | 2.06E-05 | 8 |
| SPC24 | ENSG00000161888 | -1.588922381 | 2.03E-06 | 6.77E-05 | 19 |
| XK | ENSG00000047597 | -1.588890662 | 4.89E-05 | 0.001092138 | X |
| PAPSS2 | ENSG00000198682 | -1.58489318 | 1.63E-05 | 4.27E-04 | 10 |
| CKS1B | ENSG00000173207 | -1.584654257 | 3.83E-10 | 2.83E-08 | 1 |
| RAD51AP1 | ENSG00000111247 | -1.583691422 | 4.33E-06 | 1.33E-04 | 12 |
| BUB1 | ENSG00000169679 | -1.581058027 | 1.32E-09 | 8.88E-08 | 2 |
| SPAG5 | ENSG00000076382 | -1.579377402 | 7.12E-26 | 4.08E-23 | 17 |
| KIF11 | ENSG00000138160 | -1.577176615 | 4.08E-05 | 9.37E-04 | 10 |
| APOBEC3B | ENSG00000179750 | -1.573620845 | 4.84E-15 | 8.08E-13 | 22 |
| TYMS | ENSG00000176890 | -1.571127889 | 1.46E-23 | 5.98E-21 | 18 |
| WNT10B | ENSG00000169884 | -1.570569051 | 5.25E-06 | 1.58E-04 | 12 |
| TICRR | ENSG00000140534 | -1.569392437 | 1.09E-12 | 1.29E-10 | 15 |
| SGO1 | ENSG00000129810 | -1.565896723 | 4.90E-10 | 3.51E-08 | 3 |
| GINS1 | ENSG00000101003 | -1.565069281 | 5.62E-11 | 4.79E-09 | 20 |
| | ENSG00000278238 | -1.564743875 | 9.59E-06 | 2.69E-04 | 13 |
| CDC45 | ENSG00000093009 | -1.563014627 | 2.08E-18 | 5.04E-16 | 22 |
| ATAD5 | ENSG00000176208 | -1.562128291 | 0.001398147 | 0.017907515 | 17 |
| | ENSG00000272512 | -1.561240713 | 2.54E-05 | 6.21E-04 | 1 |
| GINS4 | ENSG00000147536 | -1.559798696 | 1.69E-09 | 1.10E-07 | 8 |
| GINS2 | ENSG00000131153 | -1.559266486 | 4.49E-09 | 2.72E-07 | 16 |
| ASIC1 | ENSG00000110881 | -1.559021535 | 5.15E-11 | 4.43E-09 | 12 |
| INHBA | ENSG00000122641 | -1.555857542 | 1.45E-04 | 0.002767255 | 7 |
| FAM72D | ENSG00000215784 | -1.551876514 | 3.49E-05 | 8.21E-04 | 1 |
| ODC1 | ENSG00000115758 | -1.550287628 | 5.08E-29 | 5.15E-26 | 2 |
| PSRC1 | ENSG00000134222 | -1.54943867 | 1.73E-12 | 1.94E-10 | 1 |
